# Unsupervised learning of DNA sequence features using a convolutional restricted Boltzmann machine

**DOI:** 10.1101/183095

**Authors:** Wolfgang Kopp, Roman Schulte-Sasse

## Abstract

Transcription factors (TFs) are important contributors to gene regulation. They specifically bind to short DNA stretches known as transcription factor binding sites (TFBSs), which are contained in regulatory regions (e.g. promoters), and thereby influence a target gene’s expression level. Computational biology has contributed substantially to understanding regulatory regions by developing numerous tools, including for discovering *de novo* motif. While those tools primarily focus on determining and studying TFBSs, the surrounding sequence context is often given less attention. In this paper, we attempt to fill this gap by adopting a so-called *convolutional restricted Boltzmann machine* (cRBM) that captures redundant features from the DNA sequences. The model uses an unsupervised learning approach to derive a rich, yet interpretable, description of the entire sequence context. We evaluated the cRBM on a range of publicly available ChIP-seq peak regions and investigated its capability to summarize heterogeneous sets of regulatory sequences in comparison with MEME-Chip, a popular motif discovery tool. In summary, our method yields a considerably more accurate description of the sequence composition than MEME-Chip, providing both a summary of strong TF motifs as well as subtle low-complexity features.

## Background

Transcription factors play a fundamental role in gene regulation by binding to specific DNA sequences, termed transcription factor binding sites (TF-BSs). Thereby, they drive cell-type specific and developmental processes. TF binding is usually determined via a ChIP-seq assay. Once the ChIP-seq-derived regulatory sequences have been acquired, a multitude of tools can be employed to further investigate their content, among which *de novo* motif discovery tools have proven to be particularly helpful (e.g. MEME-Chip and RSAT [1, 2]). These tools derives a concise description of the TFBSs, usually represented as position frequency matrices (PFMs) or position specific score matrices (PSSMs), which are referred to as TF motifs. Other regulatory sequence tools, e.g. contained in the MEME suite [3] or RSAT [2], can then be used to compare similarities between TF motifs, scan for motif matches and motif enrichment with *de novo* or known TF motifs from databases, including Jaspar [4], Transfac [5], Hocomoco [6]. This eventually enables functional annotation of the regulatory sequences and potentially reveals novel players in the regulatory landscape (e.g. co-factor motifs).

While, *de novo* motif discovery tools focus on identifying TF binding motifs, they treat the remainder of the sequence as belonging to the set of background sequences, which is largely deemed to be less interesting. However, TF binding is not only characterized by the presence of TF motif matches, but by the entire sequence context. For instance, TFBSs might be embedded in CpG islands or co-localize with TFBSs of co-factors [7]. This provides motivation for the collection of a comprehensive summary of the entire sequence context, rather than merely focusing on strong TF binding sites.

Recently, deep learning, which has been successfully used in a range of fields, has also revolutionized computational biology [8, 9, 10]. Drawing inspiration from computer vision, deep learning has proven highly effective in automatically revealing DNA sequence features from raw DNA sequences that would be predictive of e.g. chromatin accessibility [10] or protein binding [8]. These features (e.g. weight matrices) recapitulate known TF motifs, co-factor motifs and low-complexity features [10], which in turn, comprehensively describe the entire sequence context. To date, deep learning-based approaches predominantly adopt a supervised learning strategy, where the relationship between raw DNA sequences and a target label (e.g. bound or unbound by a TF) is learned. On the other hand, training labels are not always available. This creates the need for alternative unsupervised learning strategies, which capture redundant structure from the sequence alone. Such approaches have, to our knowledge, garnered less attention so far.

In our article, we present an unsupervised learning model, termed *convolutional restricted Boltzmann machine* (cRBM), designed to fill this gap. Our study drew inspiration from the work of Lee *et al.* [11] who introduced a similar model tailored towards computer vision problems. We investigated the cRBM in a case study on *Oct4*, *Mafk* and *JunD* ChIP-seq peak sequences. In summary we found that, first, the cRBM reveals plausibe DNA features, including known TF motifs. Second, the cRBM leads to a more accurate description of the sequences compared to using MEME-Chip, a popular *de novo* motif discovery tool. Third, the cRBM adequately captures sequence context in heterogeneous sequences and can in turn be used to cluster sequences based in their sequence composition.

## Results

We start by introducing the cRBM and the approximate inference procedure. Subsequently, we demonstrate the model for analysis of various ChIP-seq peak regions that were obtained from the ENCODE project [12].

### Convolutional restricted Boltzmann machine

In the following, we shall discuss a convolutional restricted Boltzmann machine for learning local features from DNA sequences composed of the nucleotide letters 𝒜 = {*A, C, G, T*}, inspired by [11, 13]. We represent a DNA sequence of length *N* as |𝒜| × *N* matrix denoted by D = (*d_ij_*) in *one-hot encoding* [10]. That is, each column contains exactly *one* one and otherwise zeros where the row in which the one occurs uniquely corresponds to a nucleotide in 𝒜. We introduce the DNA features of length *M* bp as a the set of *K* weight matrices 𝒲 = {*W^k^ ∈ 𝓡^|𝒜|×M^*}_1≤*k*≤*K*_. The weight matrices 𝒲 play the role of a set of PSSMs [14]. However, they do not necessarily represent log-odds ratios at each position, but rather general weights (e.g. local energy contributions). We shall refer to the matrices 𝒲 as the motifs that are derived by the cRBM. Furthermore, we introduce the bias terms of the 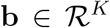 and 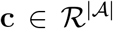. Together, 𝒲, **b** and **c** constitute the parameters of the model, which we denoted by *θ* ∈ 𝒲, **b**, **c**}. Finally, we introduce a *K* × *Ñ* matrix **H** = (*h_k,j_*) containing binary variables *h_k,j_* ∈ {0,1} for each element which indicates sequence matches of the *K* weight matrices at each position throughout the sequence **D**, where *Ñ* = *N* − *M* + 1. For example, *h_k,i_ = 1* denotes a match of weight matrix *k* at position *i* in the given DNA sequence, which we shall also refer to as a motif match.

We define the *convolutional restricted Boltzmann machine* in terms of the energy function for a joint assignment (*D*, *H*) given by

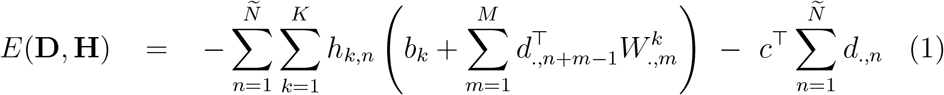

similarly as described in Lee *et al.* [11]. From the energy representation we obtain the joint distribution

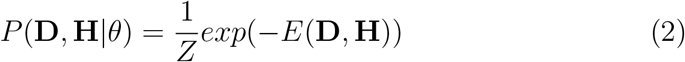

where *Z* = Σ_d,h_*exp*(—*E*(*d*,*h*)) denotes the partition function. Furthermore, the likelihood of observing a DNA sequence **D** is obtained by summing over all possible assignments of **H** according to

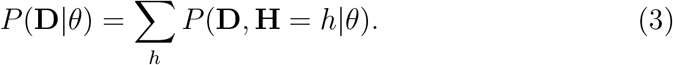

### Extension for scanning both DNA strands

DNA sequences usually occur as double-stranded hybrids in living cells which can be read with respect to the forward or the reverse strand. Hence, DNA sequence features (e.g. transcription factor binding sites) might occur on either strand equally likely.

To account for that fact, we next adapt the model to take advantage of complementary sequence information. To that end, we denote 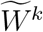as the reverse complement of *W^k^* and extend the energy function by an additional binary random variables H^z^, which represents matches of W^k^and extend the energy function by an additional binary random variables **H′**, which represents matches of 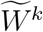.

Accordingly, the energy function becomes

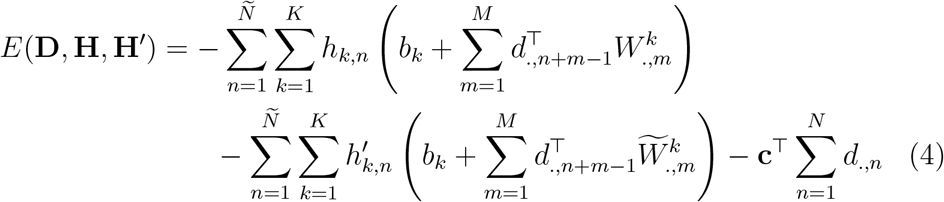

For the remainder of this article we shall focus on the cRBM that scans both DNA strands for weight matrix matches.

### Inference

Inference is a crucial ingredient in the parameter fitting procedure, which requires to compute expectations with respect to *P*(**H, H′**|D,*θ*) and *P*(**D,H,H′**|*θ*) denoted by 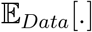 and 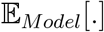, respectively. When computing these expectations, we exploit the statistical independence structure of the cRBM that renders the nucleotides at different positions in **D** conditionally independent given **H** and **H′** as well as motif matches in **H** and **H′** conditionally independent given **D**. Consequently, 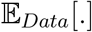 can be determined analytically. However, the analytic computation of 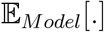 is intractable, because it requires the enumeration of exponentially many joint assignments. Therefore, we leverage a Gibbs sampling approximation to infer 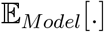 similarly as described earlier [11].

We define the following local energy contributions

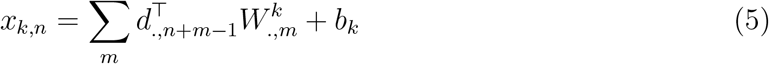

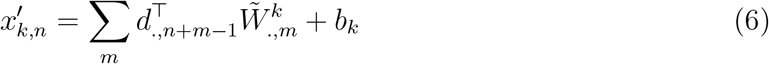

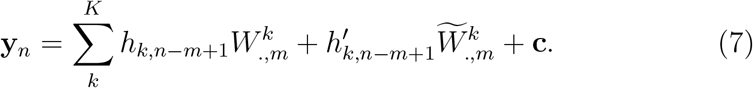

Accordingly, *x*_*k,n*_ and *x'*_*k,n*_ denote the energy contributions for observing *h_k,n_* = 1 and *H′_k,n_* = 1, given the underlying DNA sequence **D**. Moreover, *y_n_* denotes the energy contribution for observing each nucleotide at position *n* given the motif matches **H** and **H′** Note that *y_n_* represents a |𝒜|-dimensional vector.

The resulting conditional probabilities are then given by

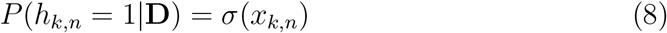

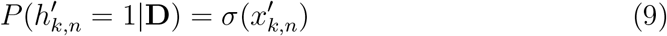

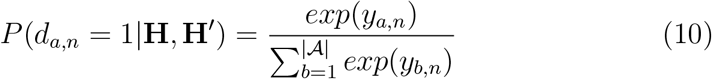

where 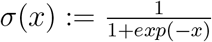.

In the Gibbs sampling procedure we take advantage of Equations (8)–(10) by alternatingly drawing samples for **H** and **H′** given **D**, and **D** given **H** and **H′**, respectively, which can be carried out in a highly parallelized fashion.

### Analysis of *Oct4* and *Mafk* ChIP-seq peaks

We applied the cRBM to ChIP-seq peak regions for *Oct4* in H1hesc cells and *Mafk* in *IMR90* cells in order to assess its capacity to learn specific and biologically interpretable DNA features in comparsion with MEME-Chip, a popular motif discovery tool.

First, we trained a cRBM with 10 motifs of length 15 bp on a set of *Oct4* and *Mafk* bound regions, separately. By inspecting the parameters, we found that both cRBMs have indeed revealed motifs that are highly reminiscent of their respective known Jaspar motifs (see Figure 1a–d). In addition, these motifs are positionally enriched in the peak center which indicates their biological relevance (see Figures S3–S4).

**Figure 1:**
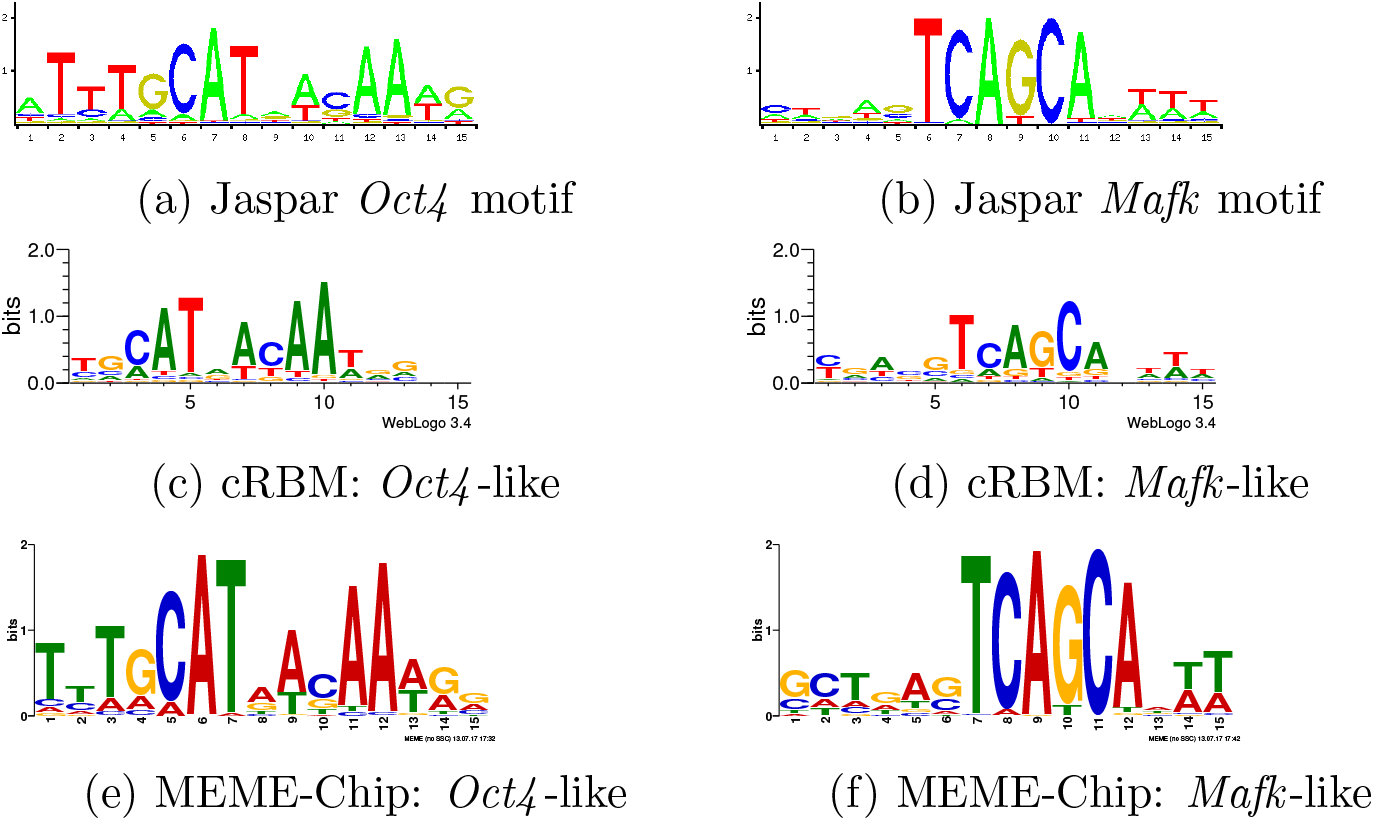
Motif comparison. a)-b) show the known Jaspar motifs for *Oct4* and *Mafk*. In comparison, c)-d) show cRBM motifs and e)-f) show MEME-Chip motifs discovered in the respective peaks which resemble the known motifs. The complete collection of revealed motifs is illustrated in Figures S1–S2 and S5–S6 for cRBM and MEME-Chip, respectively.

Besides revealing the known binding motifs for *Oct4* and *Mafk*, the cRBM has also captured several low complexity motifs (see Figures S1–S2). We sought to test whether these low-complexity motifs were just trivial common features or whether they would additionally characterize *Oct4* and *Mafk* peak regions. To that end, we used the motif abundances of the cRBM-derived motifs as input for a logistic regression classifier that attempts to discriminate *Oct4* and *Mafk* peaks (see Methods). Using all motifs extracted by both cRBMs we obtain an area under the receiver operator characteristics curve (auROC) of 0.966 (see Figure S7). By comparison, when considering only the respective *Oct4* and the *Mafk*-like motifs for the classification (see Figure 1c, S1f, S2d and 1d, respectively), the performance markedly declined to an auROC of 0.92 (see Figure S7). This indicates that even though the *Oct4* and *Mafk* motifs are the most important sequence determinants, the remaining motifs contain additional specific information about the sequence context.

Next, compared the cRBM with MEME-Chip, which is a popular tool to discover *de novo* motifs from ChIP-seq peaks. Specifically, we asked how well MEME-Chip derived motifs would represent the peaks. For that purpose, we applied MEME-Chip on the *Oct4* and *Mafk* sequences, separately, which as expected, revealed an *Oct4* and *Mafk*-like motif as well (see Figure 1e-f). Subsequently, we used all discovered MEME-Chip motifs (see Figure S5–S6) for a discrimination analysis, which resulted in an auROC of 0.886 (see Figure S7). We notice that this result is comparable with the auROC achieved by the reduced cRBM where only *Oct4* and *Mafk*-like motifs are considered for the classification. Even though a performance gap is evident, this might be partially explained by the fact that the cRBM has revealed two *Oct4* and *Mafk*-like motifs in each dataset (see Figure S1c,f and S2d,j), respectively, which were both used for the discrimination task, whereas MEME-Chip only revealed a single strong motif in each dataset (see Figure S5a and S6a). These findings indicate that apart from the discovered *Oct4* and *Mafk*-like motifs, the remaining MEME-Chip-discovered motifs only add a little extra information to characterize the sequence context.

So far we have applied the cRBM separately to *Oct4* and *Mafk* peaks, in which case the sequences are relatively homogeneous with respect to their sequence composition. Next, we sought to investigate the applicability of the cRBM for extracting features from a set of heterogeneous sequences. To that end, we trained a single cRBM using 10 motifs of length 15 bps on the combined *Oct4* and *Mafk* sequences. We found that again the *Oct4* and *Mafk* motifs were successfully revealed (see Figures S8b and S8i). Moreover, utilizing a similar discriminative analysis as described above with all 10 cRBM motifs (see Figure S8) an auROC of 0.941 was achieved (see Figure S2), which constitutes only a slight performance decline compared to when the cRBMs were trained on *Oct4* and *Mafk* sequences, separately. This suggests that the cRBM can successfully cope with heterogeneous sets of sequences.

By contrast, when using only abundances of the *Oct4* and *Mafk*-like motifs (see Figure S8b and S8i) for the logistic regression, the classification performance considerably decreased to an auROC of 0.882 compared to having trained two separate cRBMs (compare Figure 2 and S7), which underlines the relevance of not only the strong TF motifs for describing the sequence context, but also the remaining features for describing the sequence context.

**Figure 2:**
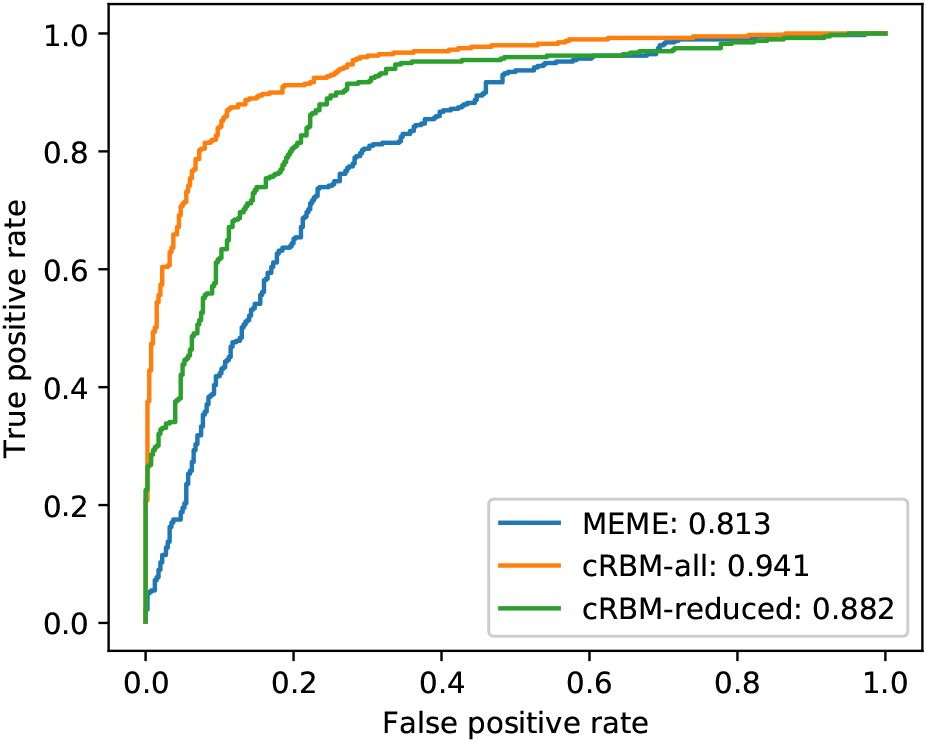
Discrimination analysis. ROC curves showing the discriminative performance of the cRBM and MEME-Chip derived motifs to discern *Oct4* and *Mafk* peaks using a logistic regression classifier (see Figure s8 and S10). For cRBM-all, all motifs were used for the classification, while for cRBM-reduced, only the *Oct4* and *Mafk*-like motifs are considered (see Figure S8b and S8i).

To put this result into perspective, we also applied MEME-Chip on the combined set of *Oct4* and *Mafk* peak sequences, which again successfully revealed an *Oct4* and a *Mafk*-like motif (see Figure S10a-c). Nevertheless, the discrimination performance based on all discovered MEME-Chip motifs (see Figure S10) substantially decreased to an auROC of 0.813 in this configuration (compare Figure 2 and S11), suggesting that the increased sequence heterogeneity considerably influences the results obtained with MEME-Chip.

Next, we examined the motif composition of *Oct4* and *Mafk* peaks to shed light on the specificity of the motifs. From the distribution of the motif abundances we observe a few motifs which appear to be shared between *Oct4* and *Mafk* peaks, including Motif 3, 4, 7 and 10 (see Figure 3 and S8). By contrast, the remaining motifs seem to be specifically enriched in either *Oct4* or *Mafk* peaks: On the other hand, Motif 1, 6 and 9 tend to be enriched in *Oct4* peaks, whereas Motif 2, 5 and 8 occur more frequently in *Mafk* (see Figures 3 and S8). Additionally, several motifs display a positional enrichment: Apart from Motif 2 and 9 (the *Mafk* and *Oct4*-like motifs; see S8), we also observe a slight positional enrichment in the peak center for Motif 4 (see Figure S8–S9). By contrast, Motif 1 and 7 appear to be slightly depleted at the peak center (see Figure S8–S9).

**Figure 3:**
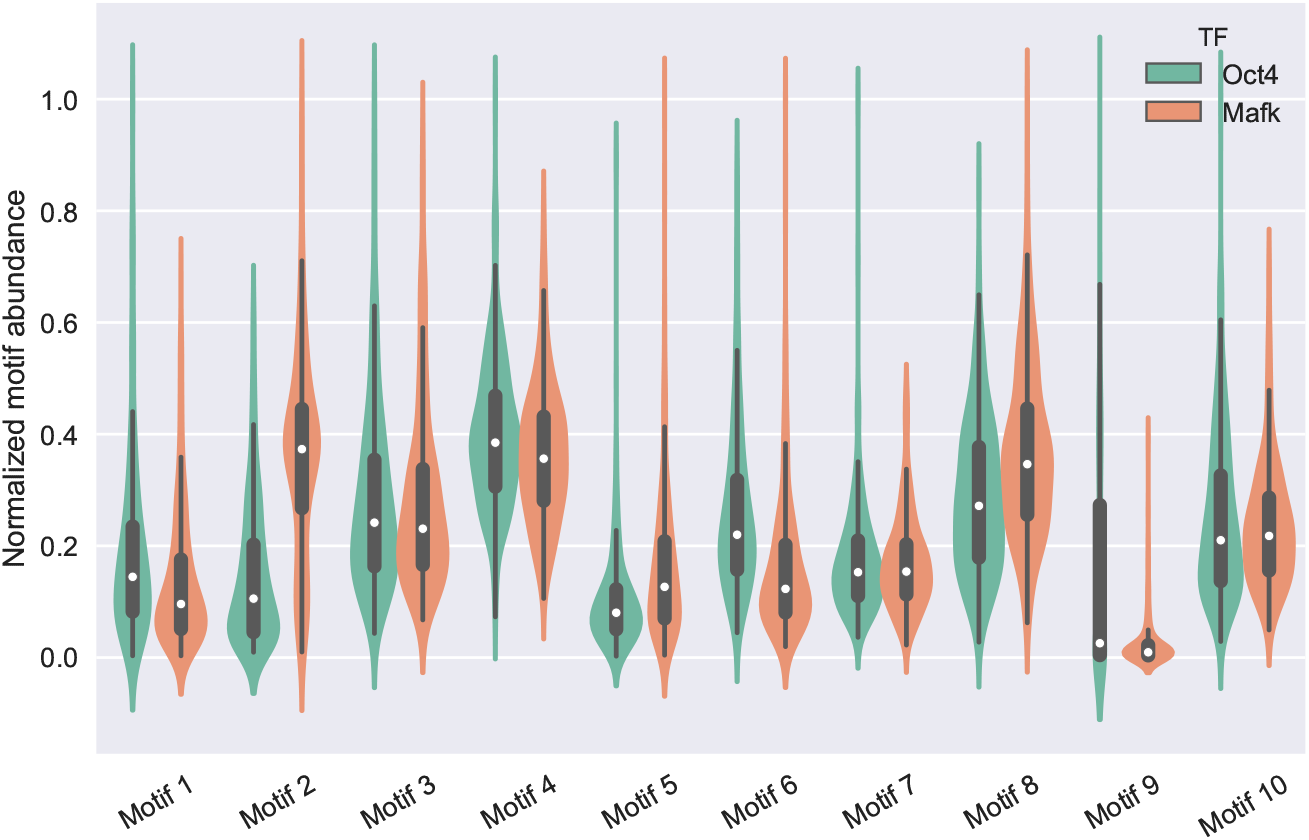
Violin plots of motif abundances. The motif abundances of motifs shown in Figure S8 are depicted in *Oct4* and *Mafk* peaks, respectively. To aid comparability, the motif abundances were normalized by the respective maximum abundance levels per motif.

In summary, this analysis suggests the cRBM yields a considerably more accurate description of the DNA sequence context compared to MEME-Chip. Not only does it identify strong known TF binding motifs, but also additional subtle features that contribute to and improve the sequence characterization.

### Discriminative analysis of *JunD* ChIP-seq peaks

In this section, we utilize the cRBM for analyzing the cell type specific binding behavior of *JunD* in *GM12878*, *HepG2*, *H1hesc*, *HeLa-S3* and *K562* cell lines, obtained from the ENCODE project [12].

We trained a single cRBM using 10 motifs of length 15 bp simultaneously on all *JunD* training peaks regardless of the cell types. A comparison with Jaspar [4] confirmed that the cRBM has successfully revealed *JunD*-like motifs (compare Figure S11 and S12i-j). Likewise, MEME-Chip unravels similar *JunD*-like motifs (see Figure S13a–b).

To further study the plausibility of the obtained cRBM and MEME-Chip motifs, we next conducted a discriminative analysis that uses the respective motif abundances as input for a multi-class logistic regression classifier to predict the cell type labels. While, the motif abundances from MEME-Chip motifs resulted in an accuracy of 40.4%, the cRBM gave rise to an accuracy of 44.9% across all cell types on the test sequences (see Table 1), which constitutes a clear improvement of sequence context explained (more than 10%) by the cRBM over MEME-Chip. Next, we explored whether and to what extent increasing the number of motifs improves the predictive performance. Accordingly, we allowed the cRBM and MEME-Chip to extract 30 motifs of length 15 bp, respectively. As expected, both methods improved in this situation compared to when only 10 motifs are used. Still, the cRBM experiences a slightly more pronounced improvement (accuracy of 47.4%; see Table 1) compared to MEME-Chip (accuracy of 42.2%; see Table 1). Moreover, MEME-Chip (using 30 motifs) still does not exceed the predictive performance achieved by cRBM with 10 motifs.

**Table 1:** *JunD* cell type specific binding prediction accuracy. This table shows the overall accuracies for predicting cell type specific *JunD* peaks based on cRBM and MEME-Chip derived motifs.

| Method | Number of motifs | Accuracy |
| --- | --- | --- |
| cRBM | 10 | 0.449 |
| cRBM | 30 | 0.474 |
| MEME-Chip | 10 | 0.404 |
| MEME-Chip | 30 | 0.422 |

Subsequently, we examined the confusion matrices for the discrimination tasks, which offer a fine-grained picture of the performances for each cell-type (see Figure S14). We observe that the cRBM-derived motifs exceed or yield similar performance levels compared to the MEME-Chip-derived motifs for most cell-types (see Figure S14). With the exception of *GM12878*, the cRBM motifs give rise to similarly many or more correct predictions compared to MEME-Chip motifs when using 10 and 30 motifs (see Figure S14). While, GM12878 is more often predicted correctly with MEME-Chip compared to cRBM-motifs (116 to 64 correct predictions, respectively, see Figure S14a and S14c), MEME-Chip motifs at the same time incur a slightly higher rate of predicting other cell types to be GM12878 (63.2% to 61.0% false prediction rate, see Figures S14a and S14c). With 30 motifs, MEME-Chip motifs still result in more correctly predicted GM12878 peaks (115 to 108 correct predictions, respectively, see Figures S14d and S14b). However, in this case the discrepancy of falsely predicted GM12878 cells becomes even more pronounced (0.59% and 0.55% for MEME-Chip and cRBM, respectively, see Figures S14d and S14b). Generally we find that the false prediction rate (normalized across columns) is lower for cRBM motifs compared to MEME-Chip motifs, except for *HeLa-S3* when 30 motifs were used. In that case MEME-Chip motifs yielded a lower fraction of other cell-types to be mis-classified as *HeLa-S3* compared to cRBM motifs (65% to 68%; see Figures S14b and S14d).

### t-SNE analysis of JunD ChIP-seq peaks

We proceed by examining a potential association between sequence similar-ties of *JunD* peaks and the respective cell-types. To this end, we use t-SNE [15] on the motif abundances of the 10 cRBM derived motifs of length 15 bp (see Methods) to project the *JunD* peak sequences onto a two-dimensional plane in which similar sequences are located in close proximity. This analysis offers the following insights: First, in concordance with the discriminative analysis described above, we find that the t-SNE analysis also unravels a cell type specific binding preference of *JunD* (see Figure 4 and S15). For instance, no two cell types exhibit the same distribution of points in the t-SNE plane. Second, *JunD* bound sequences within each cell type generally do not appear to be homogeneous. We rather observe multiple sub-populations of bound regions in each cell type with respect to the sequence composition. For instance, *HepG2* peaks can be attributed to roughly two distinct regions (see top and bottom region in Figure 4a), *K562* peaks are predominantly enriched in a single group and two less pronounced groups (see center region as well as above and below, respectively, in Figure 4c) and GM12878 peaks seem to form two groups (see bottom right and center left region in Figure 4b). Third, we appreciate sub-sets of the *JunD* bound regions that are specifically associated with a single cell type, including for *HepG2*, *GM12878*, *K562* and H1hesc (see top, bottom right, center and top right region in Figures 4a, b, c and d). On the other hand, other sub-sets of the peaks seem to contain similar sequence features across cell-types, including for *HepG2*, *K562*, *H1hesc* and *HeLa-S3* (see bottom region in Figure 4a, c, d and e).

**Figure 4:**
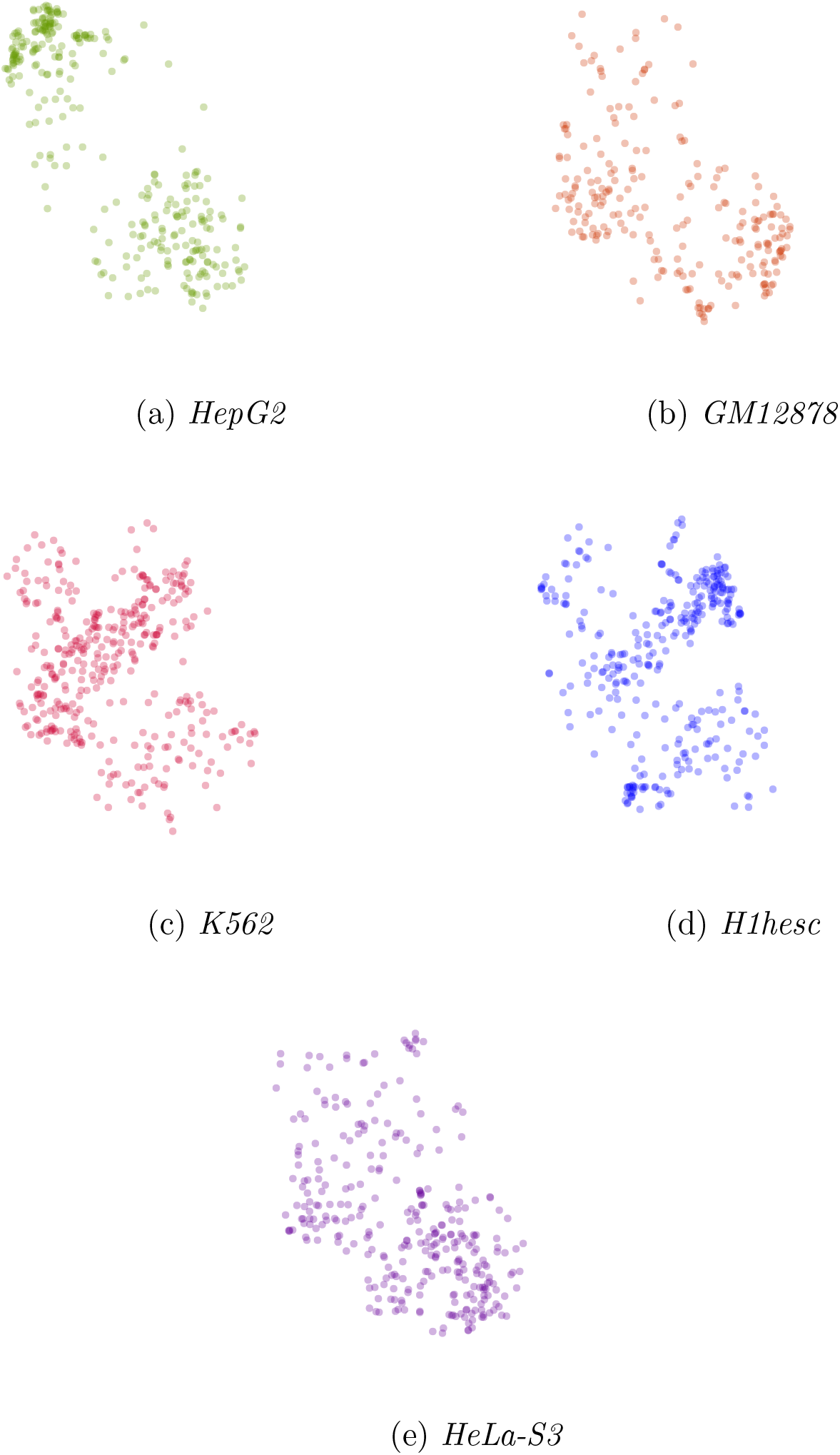
t-SNE analysis of the *JunD* peaks. Each panel shows the distribution of the cell type specific *JunD* peaks on the same t-SNE hyperplane.

This analysis suggests that peaks derived from different cell-types or conditions, but also within the same condition, seem to reflect a variety of binding modes of a TF under study. We continue by further shedding light on the different binding modes that are reflected by the differences in sequence features. For that purpose, we augment the t-SNE scatter plot (see Figure S15) with the motif abundances seen in each individual sequence (see Methods).

On a course-grain level, we appreciate three sub-populations of peaks across all cell-types: 1) peaks exhibiting the canonical *JunD* motif (see lower region in Figure 5 and Figure S12i), 2) peaks exhibiting an alternative *JunD* motif (see top region in Figure 5 and Figure S12j) and 3) peaks which lack a strong *JunD* binding motif (see center region in Figure 5).

**Figure 5:**
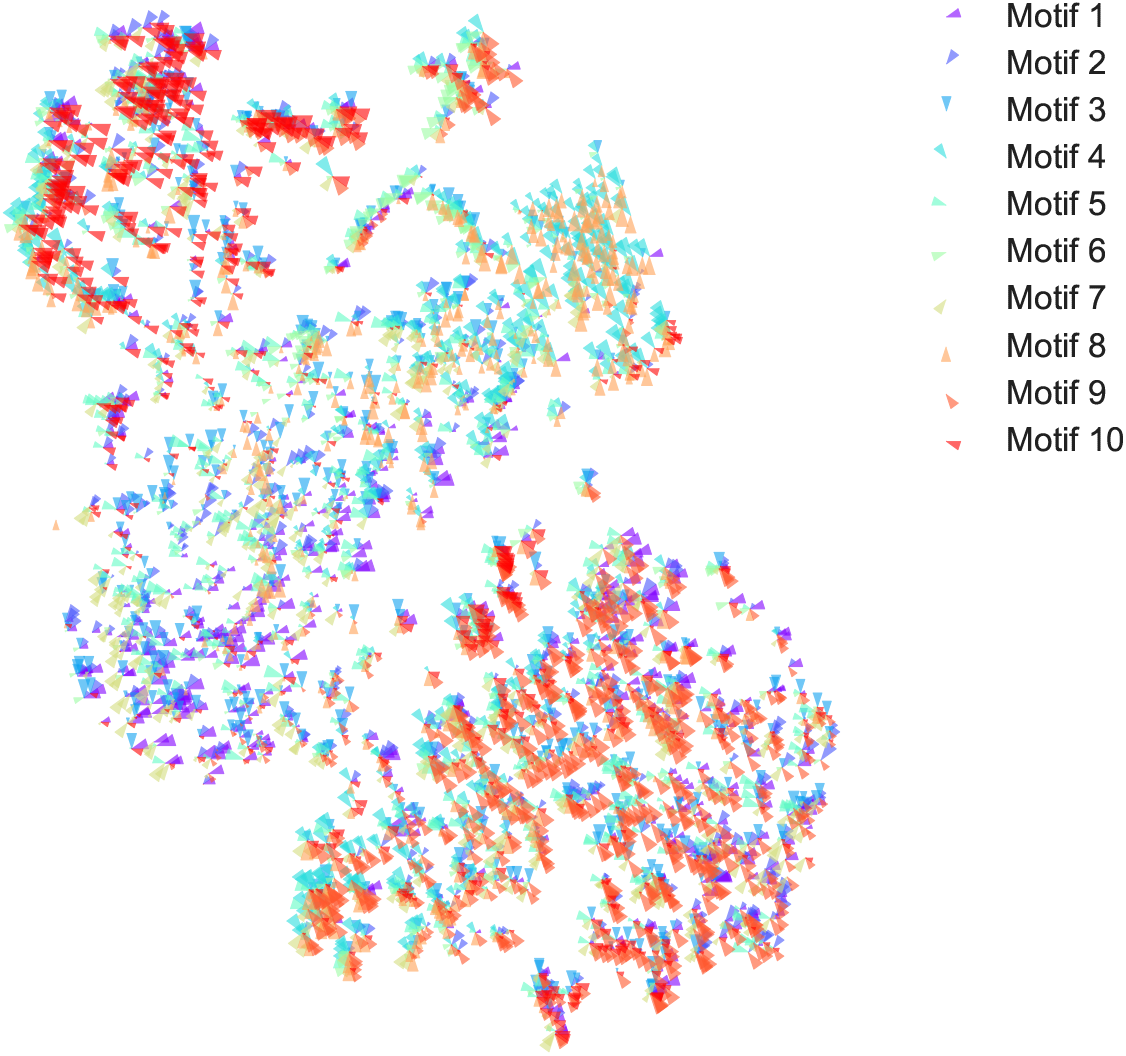
Sequence composition of ***JunD*** peaks. Using the same t-SNE hyper-plane as shown in Figure 4, each sequence is represented as a pie cart with the pie pieces corresponding to motifs (see Figure S12). The sizes of the pie pieces reflect the motif abundances.

Peaks represented by the lack of *JunD* binding motifs seem to be characterized by a number of low-complexity motifs: On the one hand, G-rich motifs (Motif 4 and 8; see Figure S12d and h) seems to appear abundantly in H1hesc peaks (compare Figure 4d and 5). On the other hand, T - and C-repeat motifs (Motif 1 and 3; see Figure S12a and c) seem to frequently occur in a subset of *K562*, *GM12878* and *HeLa*-S3 peaks (see Figure 4b,c,e and 5). These peaks could potentially reflect indirect *JunD* binding together with other TFs.

It appears that the regime characterized by the the canonical motif variant of *JunD* is shared across multiple cell types (compare on bottom regions of Figures 4 and 5), suggesting a common binding mode of *JunD* across the cell-types. On the contrary, the peaks characterized by the alternative *JunD* binding motif (see Figure S12j) seem to be specific for *HepG2*, suggesting an alternative binding mode of *JunD* in that cell line. This hypothesis is further supported by Srivastava *et al.* [16], who have reported different binding modes of *JunD* in rat: Accordingly, the 7-mer motif (TGAGTCA) is associated with homo-dimer binding, whereas, the 8-mer variant (TGACGTCA) is associated with hetero-dimer binding in conjunction with an ATF-family member.

To summarize, the cRBM has revealed an accurate and representative set of DNA features of the *JunD* bound regions, including known TF motifs, CG-rich and AT-rich features. We have shown that the cRBM yields a more comprehensive description of the sequences compared to MEME-Chip. We have further illustrated the utility of the cRBM motifs for a t-SNE analysis, which has revealed cell-type specific sequence signatures of *JunD* peaks and heterogeneous sets of peaks within each cell-type. By projecting the sequence composition on the the t-SNE plane two binding modes of *JunD* were revealed in *HepG2*, represented by occurrences of distinct *JunD* motif occurrences.

## Discussion

In this article, we presented a novel unsupervised model to extract redundant DNA sequence features from a given set of sequences. The method is termed convolutional restricted Boltzmann machine and was inspired by Lee *et al.* [11]. It was designed to take features on both DNA strands into account to acquire a comprehensive summary of the sequence composition of the regulatory sequence under study, rather than focusing only on strong TF motifs.

We have compared the cRBM with the popular state-of-the-art motif discovery tool MEME-Chip, on a number of ChIP-seq peak regions obtained from the ENCODE project [12]. Our results support the conclusion that the cRBM yields a more comprehensive and accurate summary of the regulatory sequences under study. While both methods generally agree on strong TF binding motifs in the respective ChIP-seq peak regions, the conventional motif discovery approach attributes the remaining sequence context to a background model, which is deemed less important [1, 2]. On the other hand, the cRBM extracted additional sequence features that distinctly improve the characterization of the regulatory regions.

Furthermore, our results support the conclusion that the cRBM is more accurate for summarizing heterogeneous sets of sequences than the conventional motif discovery tool. This aspect is of particular importance, as our analysis, in concordance with earlier publications [17, 18], suggests that TFs might enact different binding modes at different genomic loci even within the same condition (e.g. *JunD* binding in homo - and hetero-dimer mode in *HepG2*). This in turn is reflected by a different sequence composition for each mode and therefore a potential increase in sequence heterogeneity. The cRBM can aid the dissection of these heterogeneous sets of sequences into fairly homogeneous sub-populations which are potentially associated with different regulatory roles [17].

In our *JunD* case study, we found that a subset of binding sites lack the presence of the primary motifs (e.g. *JunD* peaks without the *JunD* motif), which potentially correspond to indirect binding events of *JunD*. While, we found that these regions can be characterized by low-complexity motifs, the question still remains as to why no additional co-factor binding motifs were revealed in that regime. One reason for this might be that the co-factor motifs occur too rarely for the cRBM (but also for MEME-Chip) to reach its detection limit.

A potential drawbacks of the cRBM is the number of hyper-parameters that need to be chosen for model fitting, which includes the learning rate α, batch size and motif hit sparsity ρ, etc. It is difficult to suggest a generally applicable set of hyper-parameters, since that choice depends on the dataset and application. We have investigated a large number of hyper-parameters for the *Oct4* and *Mafk* dataset, from which we concluded that, first, it is beneficial to use a relatively high learning rate (e.g. α = 0.1) and relatively small batch sizes (e.g. 10-20). We believe that this adds noise to the training phase which helps to overcome poor local optima when using persistent contrastive divergence. Second, we observed that it is important to start the training off with rare motif hit occurrences (e.g. ρ = 0.01). Too many initial motif hits (e.g. ρ = 0.1) make it difficult to untangle the sequence information captured by the individual weight matrices as they would lead to overlapping motif matches. That in turn impairs the interpretability and quality of the results. On the other hand, a too few motif matches (e.g. ρ = 10^-4^) tends to significantly slow down the training progress and also leads to results of low quality. The effect of other training-related hyper-parameters on the quality of the results, e.g. the sparsity enforcement λ, appeared to be noticeable, but less pronounced.

Moreover, we intentionally tested the same set of hyper-parameters for studying different ChIP-seq peaks under different situations (homogeneous and heterogeneous sets, different sequence numbers, etc.). This also confirmed that our proposed hyper-parameters, even though they might not be optimal for every situation, tend to yield reproducible and robust results for a range of different situations.

Apart from the training-related hyper-parameter, the number of motifs and the motif lengths also need to be defined prior to the analysis. Again, it is difficult to suggest an ideal strategy for choosing these model parameters optimally for all possible purposes. Generally, we have observed that increasing the number of motifs and motif lengths tends to improve the performance in the discriminative setting, but the performance gain saturates with a increasing number of model parameters. We recommend using a small number of motifs if possible, which often yields a similar sequence summary, but has the advantage of maintaining interpretability of the results (e.g. by inspection of the sequence logos). Additionally, not only may a large number of sequence logos be difficult to analyze visually, but also may the cRBM be provided with the flexibility to express e.g. a single known TF motifs in terms of multiple very similar weight matrices. This potentially captures additional subtleties in the DNA sequences, but also introduces difficulties for the interpretation the results.

In the future, we shall investigate the potential of convolutional deep Boltzmann machines [11]. Such a model would overcome the assumption that motif matches occur independently for all sequences by learning co-occurrence patterns of motifs which might further improve the sequence model.

## Conclusion

We present a novel unsupervised method, termed a convolutional restricted Boltzmann machine, to discover redundant DNA sequences features. The method facilitates the analysis of regulatory sequences by not only extracting TF binding motifs, but to extract a rich set of features capable of accurately representing the entire sequence context. In our work, we have demonstrated the utility of the method applied to publicly available ChIP-seq datasets. We believe that our model can be a valuable tool for studying regulatory sequences.

## Methods

### Datasets

We downloaded ChIP-seq peak regions in narrowPeak format from the ENCODE project [12] for the transcription factors *Oct4* in *H1hesc*, Mark in *IMR90*, *JunD* in *GM12878*, *H1hesc*, *HeLa-S3*, *HepG2* and *K562* and retained the top 4000 most significant peaks for the analysis. Furthermore, we trimmed the peaks to be 200 bp in length centered around the peak midpoint. For *JunD*, we further removed peaks that occur in multiple cell types.

This led to a total of 3997 *Oct4* peaks, 4000 *Mafk* peaks as well as 2261, 3318, 3166, 3033 and 3714 *JunD* peaks in *GM12878*, *H1hesc*, *HeLa-S3*, *HepG2* and *K562*, respectively.

Finally, we randomly split the regions into training and test regions, consisting of 90% and 10% of the sequences, respectively. While training sequences were used to fit the models, the test sequences were only used for performance evaluation.

### Training

The parameters *θ* of the cRBM were adjusted on a given set of training sequences 𝒟 = {*D*^1^, …, *D^S^* } *using persistent contrastive divergence* (PCD) [19]. The method starts with initial parameters *θ*_0_ and iteratively refines them in small steps. The weight updates are determined by computing 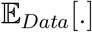 and 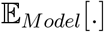 making use of the inference procedure described in the Results section.

Finally, the weights are updated on mini-batches by additionally leveraging a momentum term that damps oscillations [13].

#### Sparsity constraint

To prevent the cRBM from learning trivial features (e.g. a motif that explains the occurrence of only a single nucleotide) we employ a sparsity constraint during training, which penalizes too frequent occurrences of weight matrix matches *P*(**H|V**) (and *P*(**H′|V**)).

Therefore, we adopt the cross-entropy-based penalty term [13]

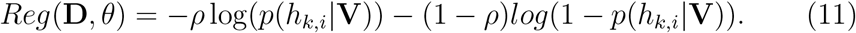

where *ρ* denotes a pre-defined, desired target frequency for weight matrix matches to occur.

During training, we incorporate the regularization term into the training ob jective according to

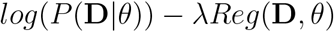

where λ denotes the *Lagrangian* multiplier which corresponds to the strength of enforcing the penalty.

#### Weight initialization

Weight initialization plays a critical role in the cRBM’s capability to extract meaningful and specific DNA features. While, the weight matrices would usually be initialized with small values sampled from a Gaussian distribution and setting the bias terms *b_k_* and **c** to zero, we noted that this leads to poor results when studying DNA sequences.

We found that it was important to initiate the bias terms *b_k_* with negative values, such that motif matches occur rarely when training is initiated (see Equations (5)–(6) and (8)–(9)). Accordingly, we implemented the following initialzation approach: We randomly initialized the weight matrices by sampling from 𝒩(0,1). Assuming that the nucleotides occur i.i.d. in a given sequence, the distribution of the aggregated weights across the weight matrix length *M* is distributed according to 𝒩(0, *M*). Under this assumption and to enforce that an initialized weight matrix only causes a match with probability *P*(*h_k,n_* = 1|**V**) = ρ, we sought to identify the associated bias term according to

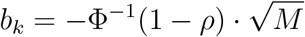

where Φ^-1^ denotes the inverse cumulative normal distribution function.

#### Training parameters for the cRBM

We trained the cRBM on *Oct4* and *Mafk* training sequences with 10 motifs of length 15 using the following training hyper-parameters: Learning rate α = 0.1, mini-batch size of 20, momentum term of 0.95, ρ = 0.01, λ = 0.1, and 300 epochs.

We used the same training hyper-parameters to train a single cRBM on the combined *Oct4* and *Mafk* sequences, as well as for the combined *JunD* sequences, to evaluate the usefulness of the hyper-parameters across different situations. Additionally, we trained a cRBM with 30 motifs on *JunD* peak regions with the same hyper-parameters.

### Motif discovery with MEME-Chip

We utilized MEME-Chip to identify the 10 best motifs of length 15 bp on *Oct4* and *Mafk* peaks separately as well as on the combination of the peaks. Similarly, we used MEME-Chip to reveal the 10 and 30 best motifs of length 15 from the combined set of *JunD* training sequences across cell types.

### Motif abundance using the MEME-Chip motifs

For all MEME-Chip-derived motifs we scanned the peak sequences using the obtained position-specific score matrices, resulting in match scores *s_i_* across all positions *i* of a given sequence. Forward and reverse strand contributions were summed. Subsequently, the motif match probabilities were determined according to 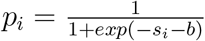 where *b* represents an offset that indicates how frequently motif matches occurs in general. Finally, the motif abundance is computed by summing up the match probabilities *p_i_* across a given sequence.

In order to ensure a fair comparison with the cRBM, we optimized the offset parameter b to achieve the best discriminative performance. To this end, for the *Oct4* and *Mafk* peak analysis, we determined the offset b for each motif individually that best separates the training sequences in terms of the auROC. For the multi-class prediction problem of the cell type specific to the *JunD* peaks, the best discrimination performance was achieved by optimizing the accuracy on the training set by adjusting a common offset b across all motifs, rather than an individual offset per motif. We found that the latter strategy resulted in a substantially poorer classification result.

### Motif abundance using the cRBM motifs

For all weight matrices, the local energy contributions for the forward and reverse strand (Equation (5)–(6)) were summed up to determine the match probability *p_i_* at position *i* in a given sequence. Subsequently, the motif abundance was obtained by summing match probabilities *p_i_* up across all positions *i*.

### Discriminative analysis

For the discrimination analysis of the peak sequences we employed logistic regression. As input the motif abundances of the cRBM or MEME-Chip motifs were used, respectively. For the case where the cRBM was used to determine motifs in *Oct4* and *Mafk* peaks, separately, the motifs of both models were merged. Likewise, we merged the motifs of the MEME-Chip analysis. The discrimination was performed as a binary classification task to predict *Oct4* and *Mafk* peak as well as a multi-class classification task to predict the cell types of origin (*GM12878*, *H1hesc*, *HepG2*, *HeLa-S3* and *K562*) for the *JunD* peaks.

For the binary classification task we evaluated the performance using the auROC, while for the multi-class prediction task we utilized the accuracy and the confusion matrix.

### t-SNE analysis of sequence similarity

We used the motif abundances obtained from the cRBM motifs on *JunD* peaks as input for the t-SNE algorithm [15], implemented in scikit-learn [20] using default parameters. Thereby, we restricted the algorithm to project the data points onto a two-dimensional hyper-plane.

Furthermore, we illustrated the motif abundances of the individual sequences in the t-SNE hyper-plane by representing each motif as a pie chart. The size of the motif was displayed in proportion to max(0, *a_k_* – median(*a_k_*)) where *a_k_* denotes the abundance level of motif *k* in a given sequence and median(*a_k_*) denotes the median abundance level observed across all sequences.

### Implementation

We have implemented the model in Python using the package Theano [21]. For the performance assessment we have employed [20]. The method is freely available on github https://github.com/schulter/crbm.

## Author’s contributions

WK conceived the method and designed the analysis. RSS implemented the cRBM with help from WK. WK carried out the analysis with help from RSS. WK and RSS wrote the article.

## Acknowledgements

The authors wish to thank Annalisa Marsico for discussions and Brian Caffrey for proofreading.

---

**Figure S1:**
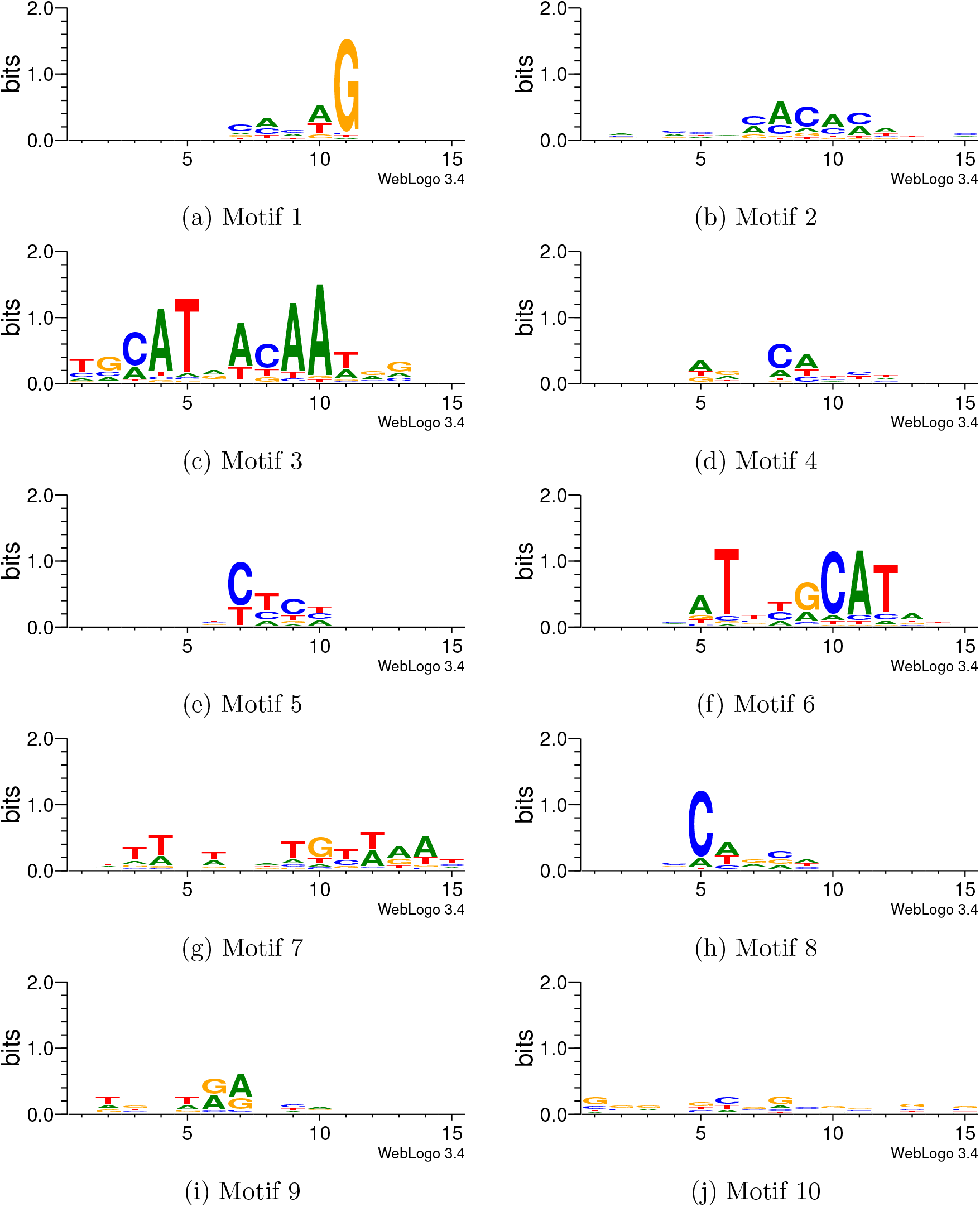
Motifs discovered by training the cRBM on *Oct4* peaks.

**Figure S2:**
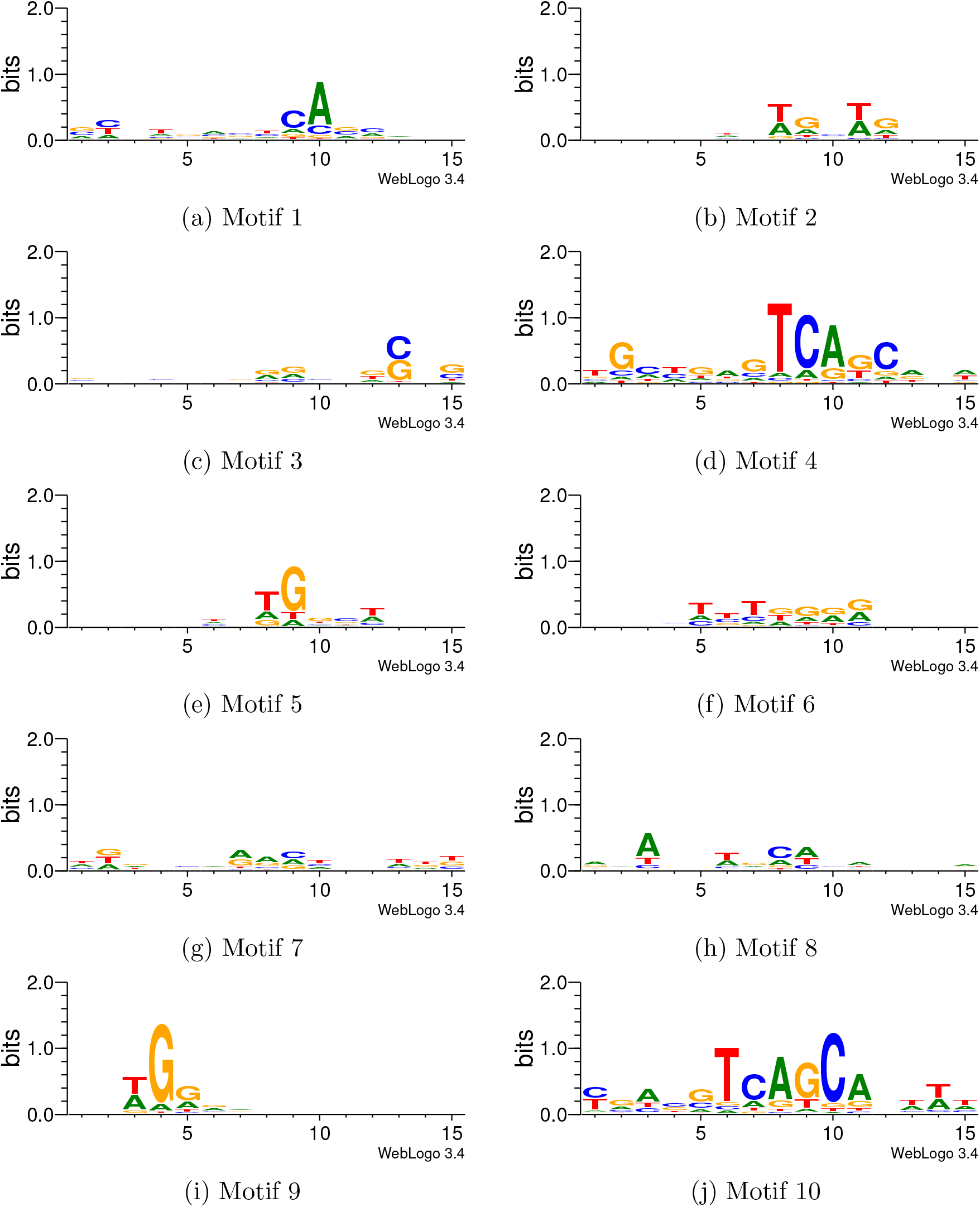
Motifs discovered by training the cRBM on *Mafk* peaks.

**Figure S3:**
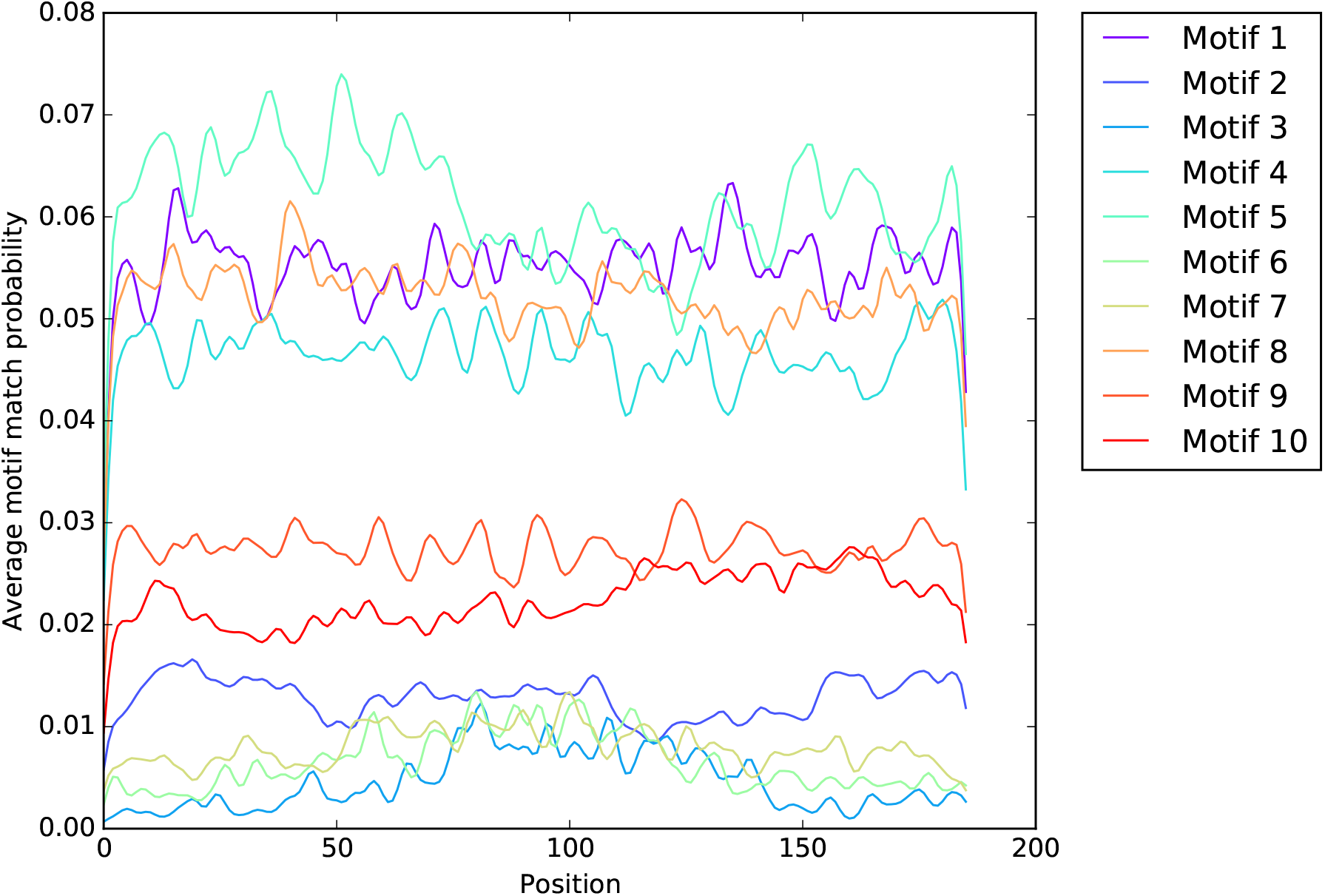
Average smoothed profile of motif matches on *Oct4* test regions using the *Oct4*- specific cRBM. Motif 3 and 5, which show a resemblance to the known *Oct4* motif (see Suppl. Figure S1), preferentially occurs at the peak center.

**Figure S4:**
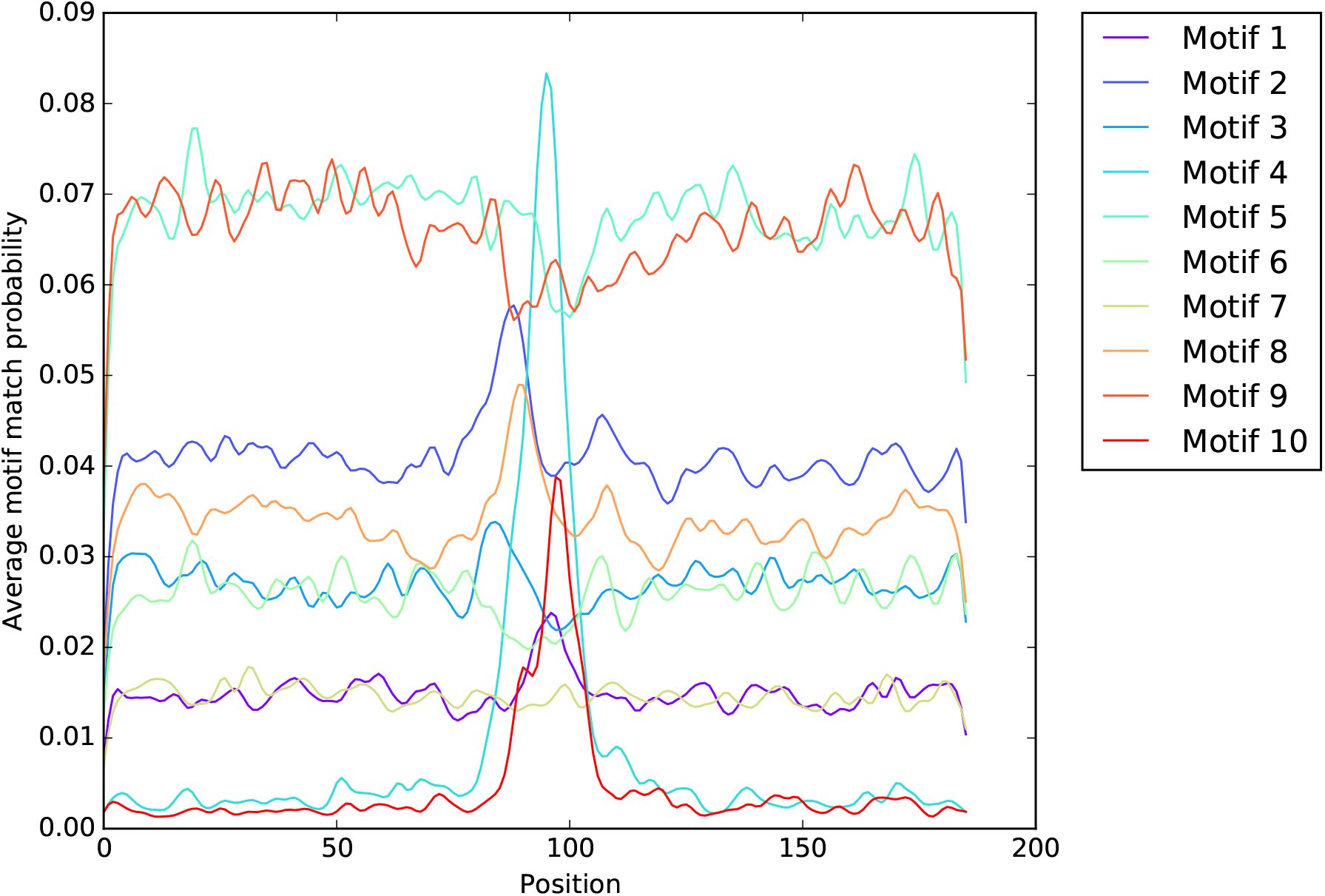
Average smoothed profile of motif matches on *Mafk* test regions using the *Mafk*- specific cRBM. Motif 4 and 10, which shows a resemblance to the known *Mafk* motif (see Suppl. Figure S2), preferentially occurs at the peak center.

**Figure S5:**
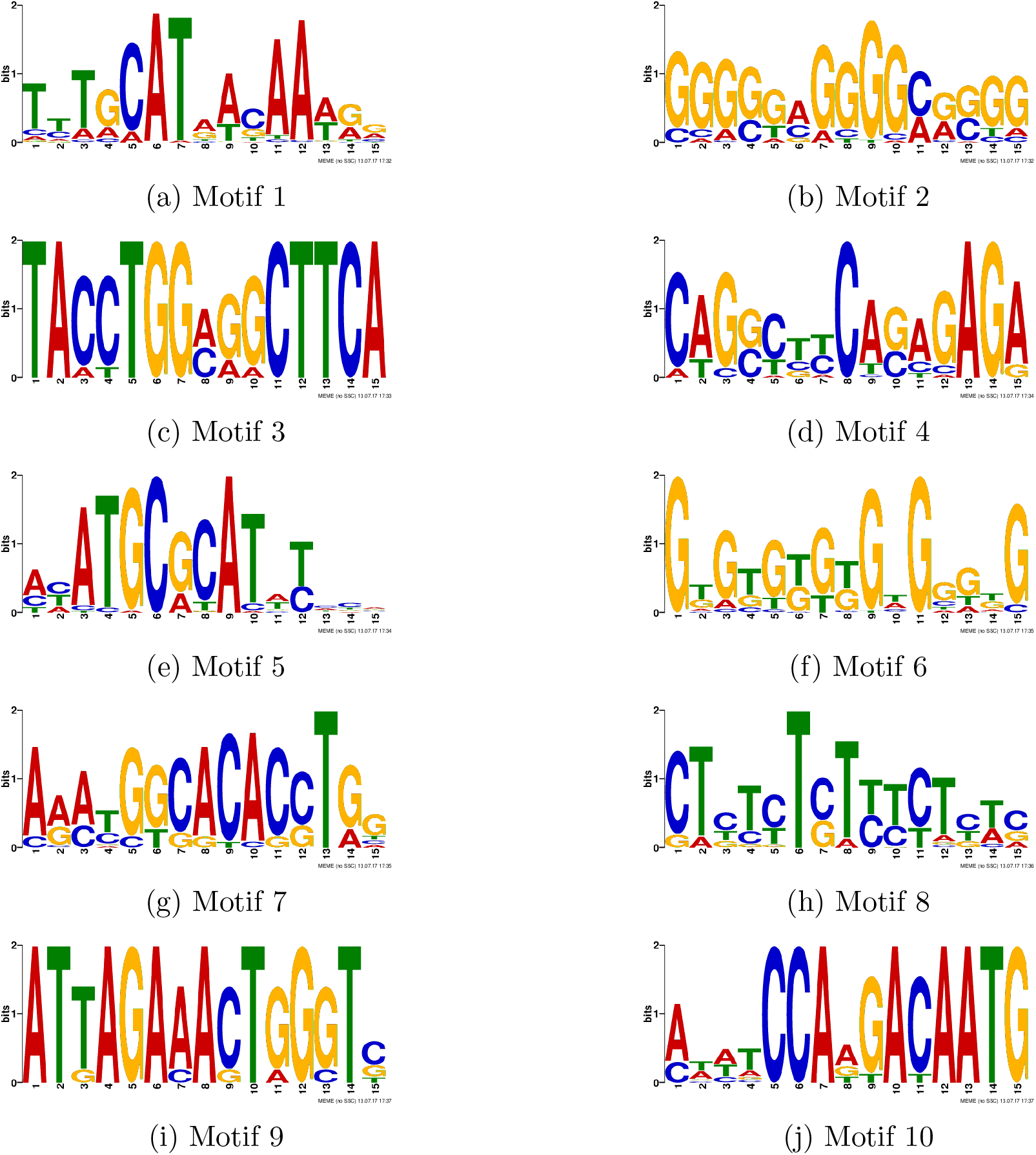
MEME-Chip discovered motifs in *Oct4* peaks.

**Figure S6:**
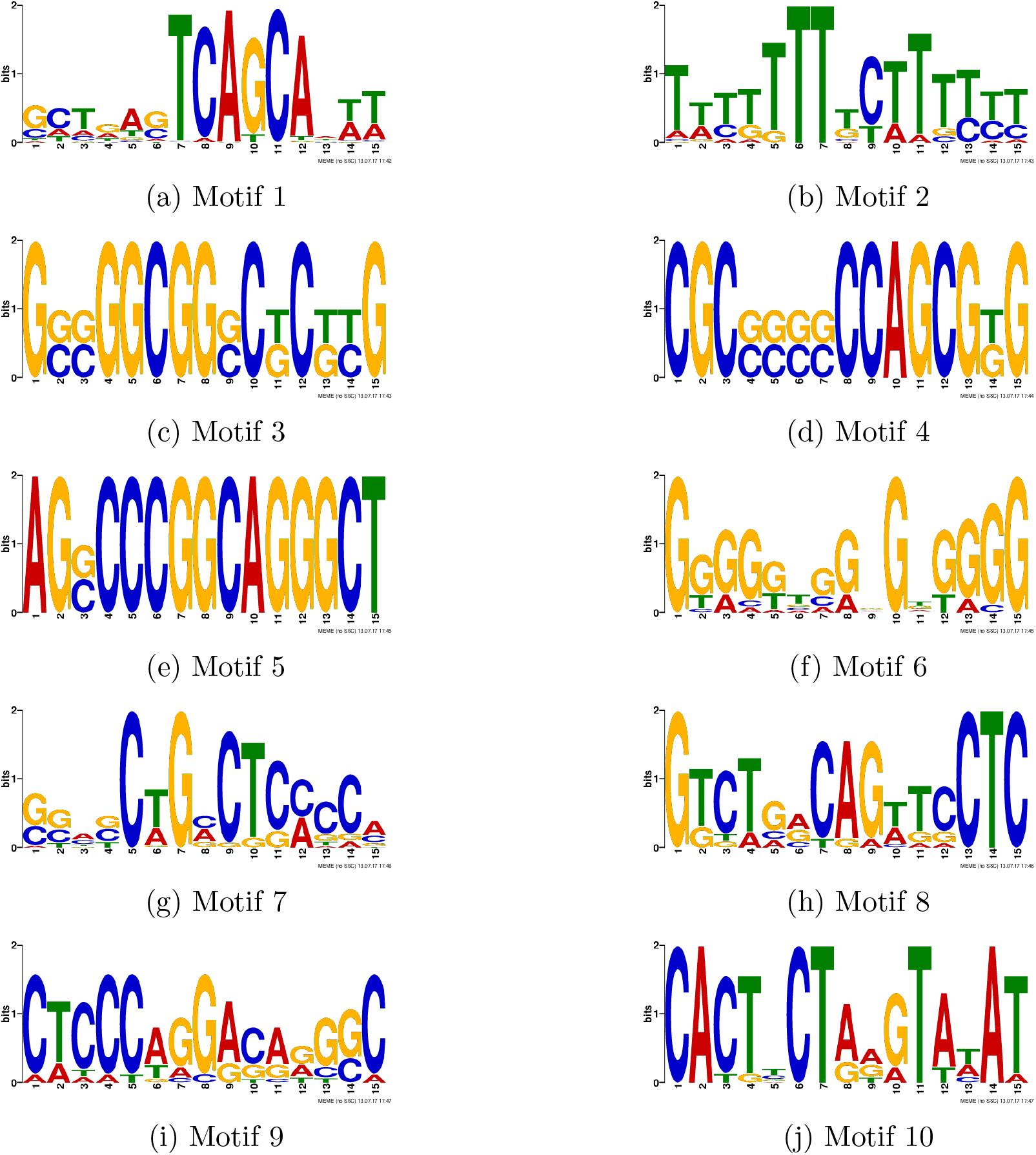
MEME-Chip discovered motifs in *Mafk* peaks.

**Figure S7:**
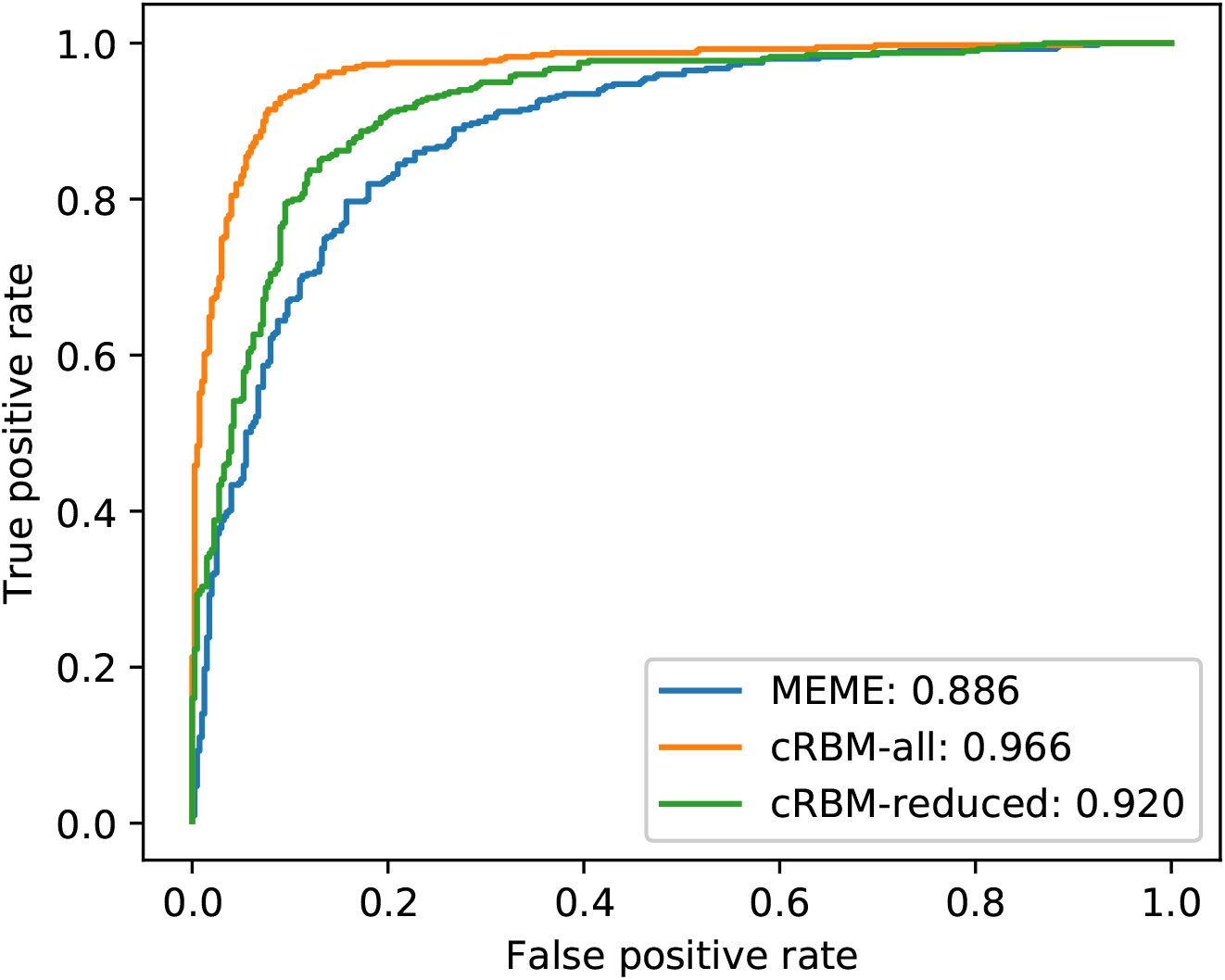
ROC curve showing the discriminative performance for classifying *Oct4* and *Mafk* test peaks. For all cases, cRBM and MEME-Chip were used to discover motifs in *Oct4* and *Mafk* peaks (see Suppl. Figure S1–S2 and S5–S6, separately. For these motifs, the abundances were provided as input for a logistic regression classifier. While for cRBM-all, all motifs where used, the cRBM-reduced prediction only relies on Motif 3 and 6 (see Suppl. Figure 1c and f) Motif 4 and 10 (see Suppl. Figure 2d and j), which resemble the known binding motifs of *Oct4* and *Mafk*.

**Figure S8:**
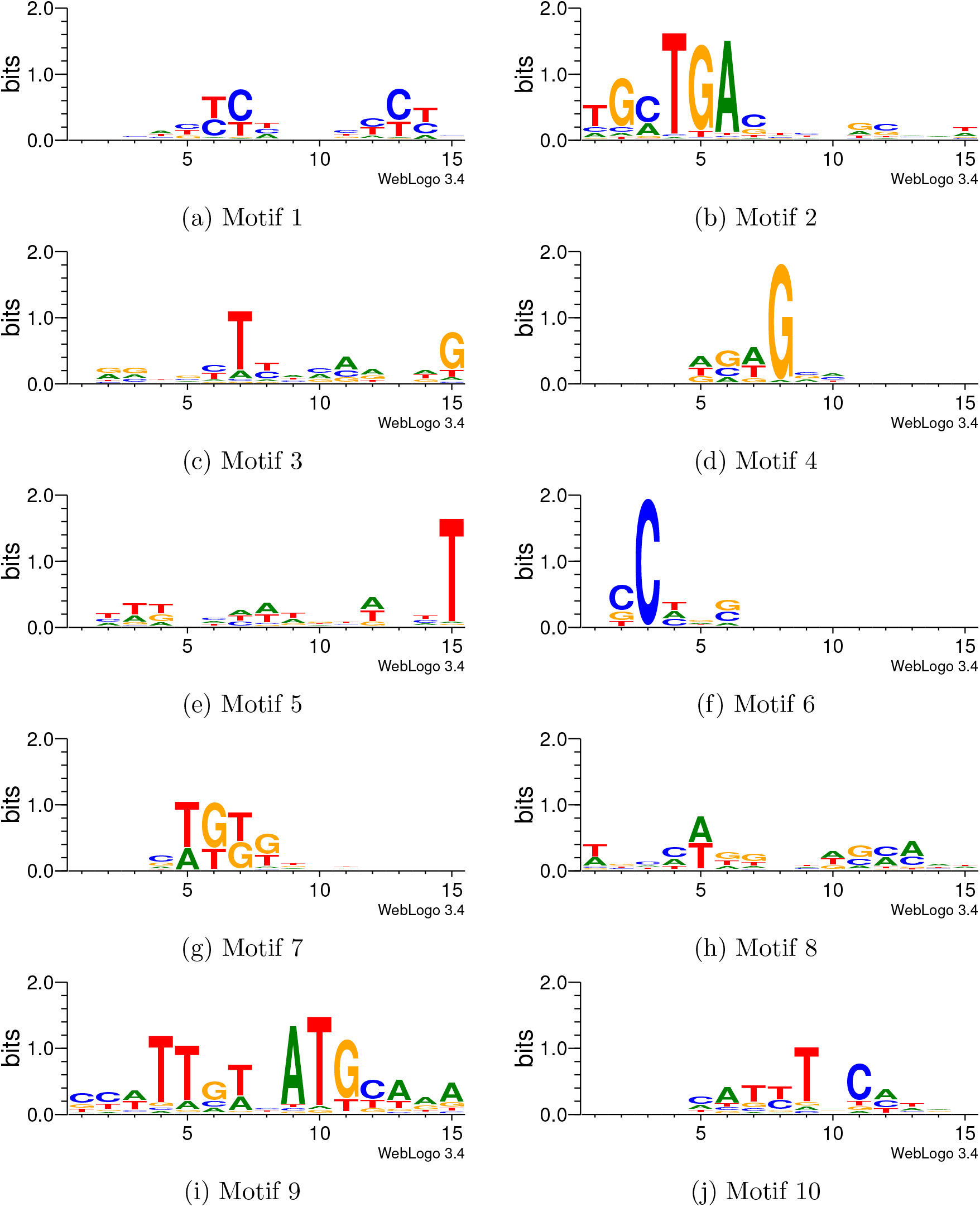
Motifs discovered by training a single cRBM on *Oct4* and *Mafk* peaks, simultaneously.

**Figure S9:**
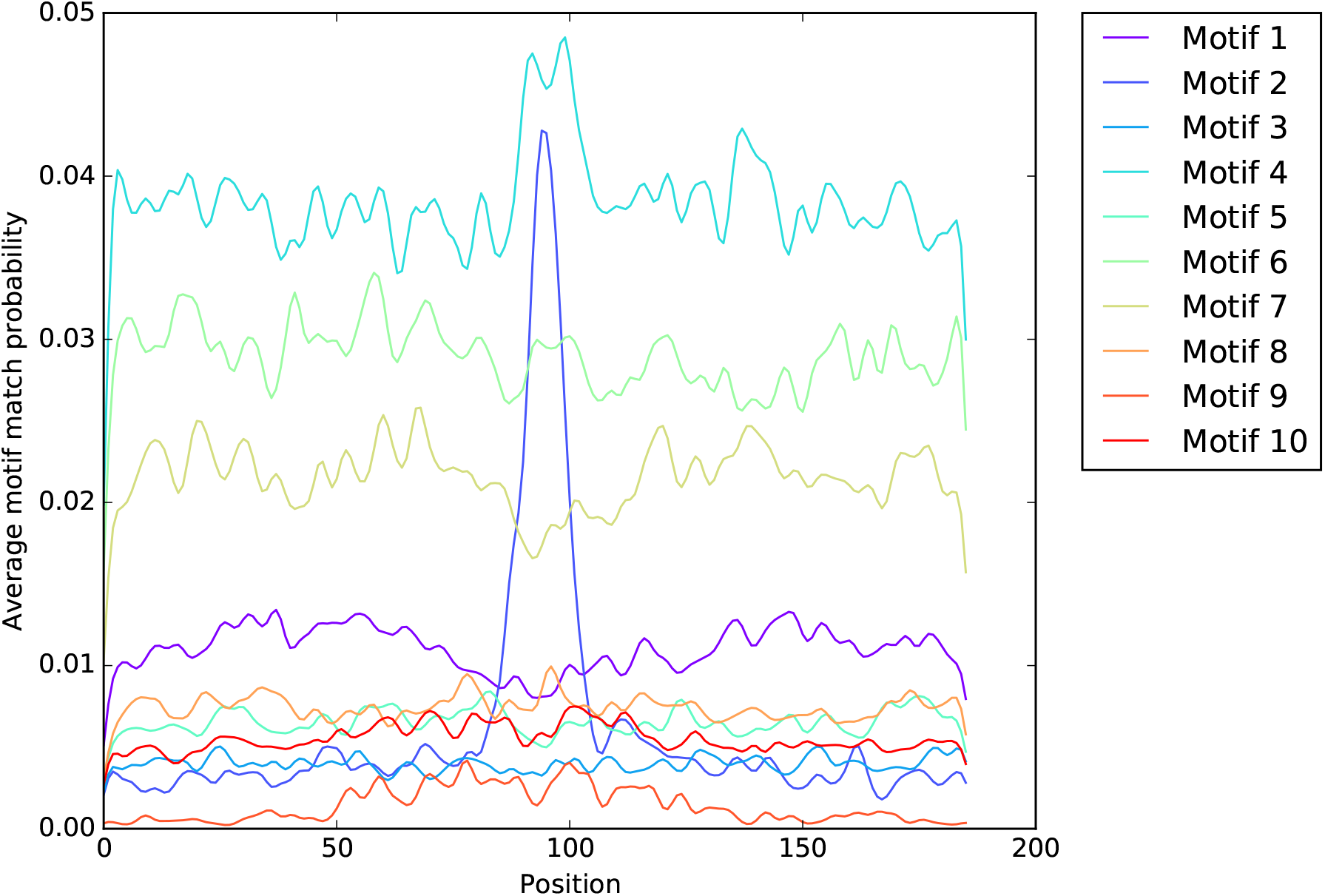
Average smoothed profile of motif matches on *Oct4* and *Mafk* test regions resulting from a single cRBM. The corresponding motifs are depicted in Suppl. Figure S8.

**Figure S10:**
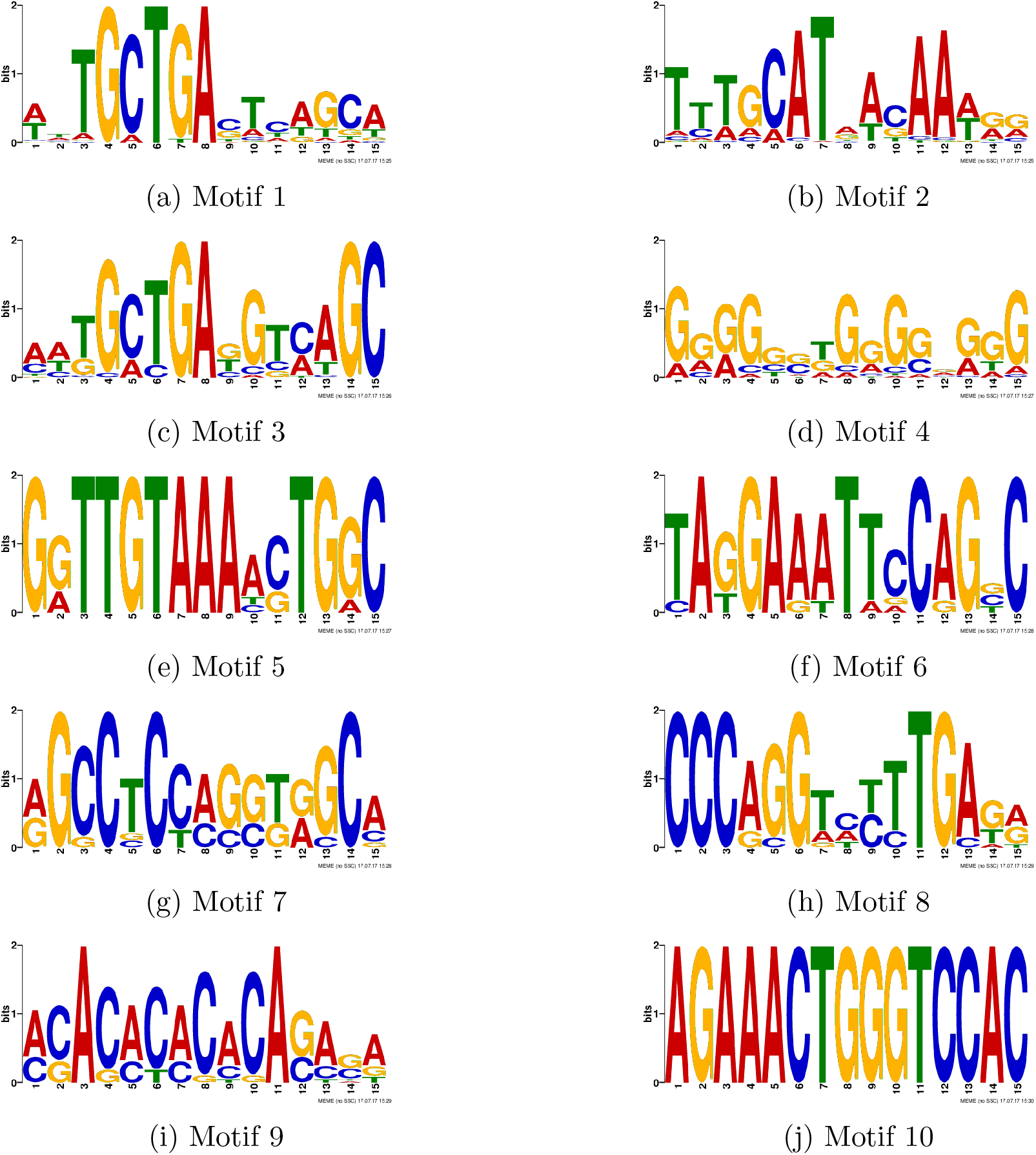
MEME-Chip discovered motifs in the combination of *Mafk* and *Oct4* peaks.

**Figure S11:**
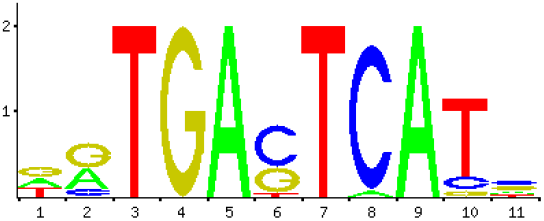
Jaspar *JunD* motif.

**Figure S12:**
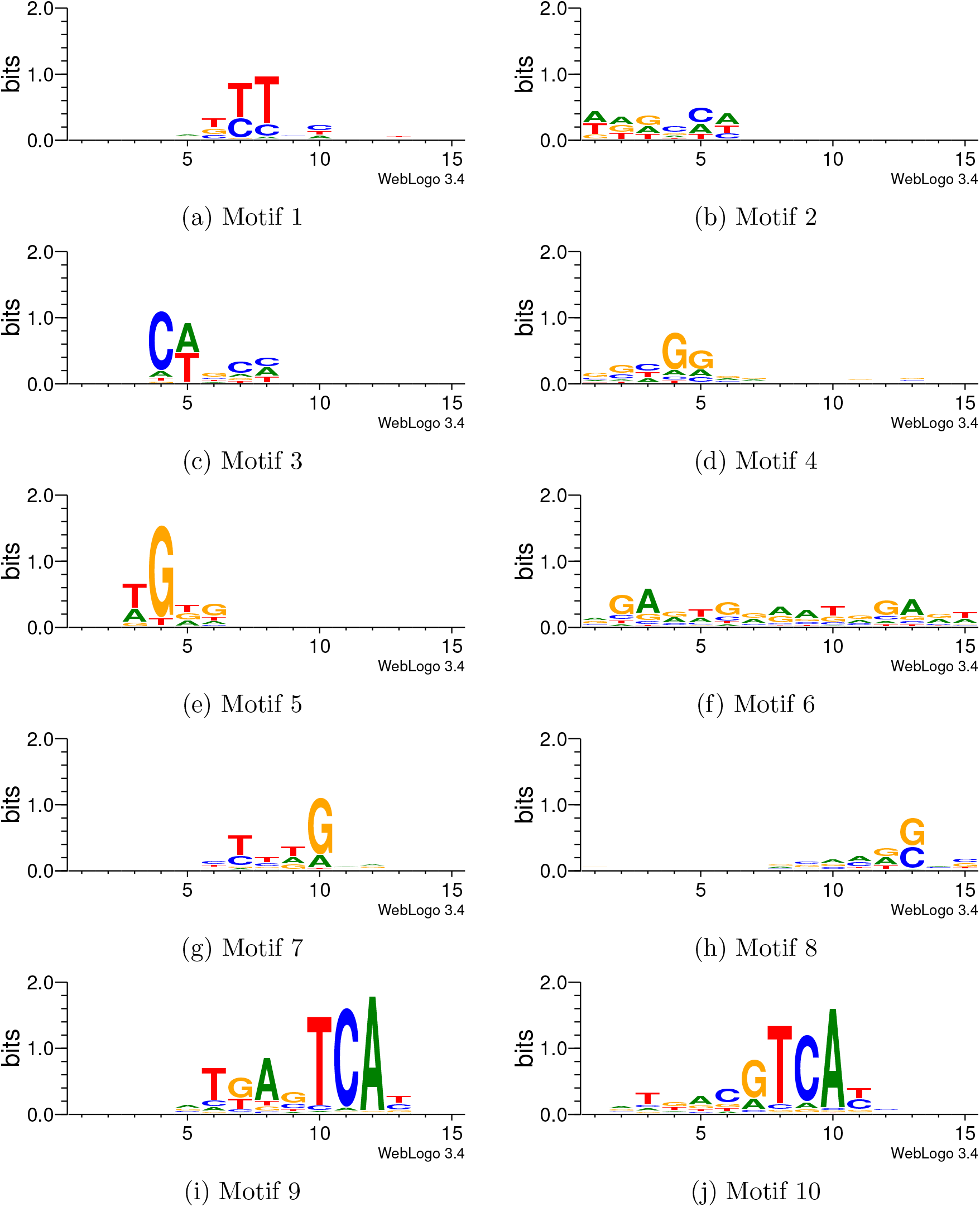
Motifs discovered by training the cRBM in *JunD* peaks across multiple cell types.

**Figure S13:**
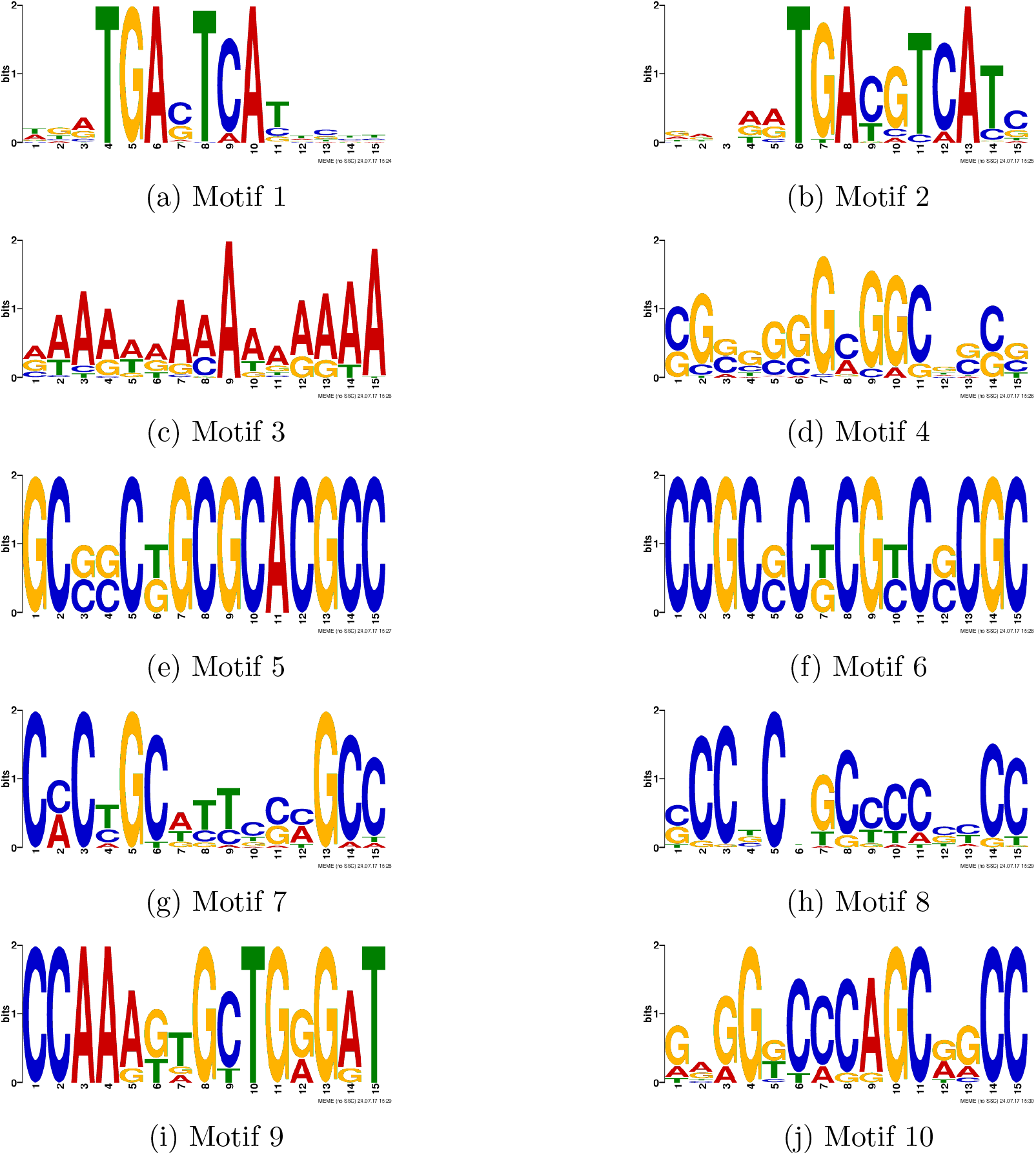
MEME-Chip discovered motifs in all *JunD* peaks regardless of cell-type.

**Figure S14:**
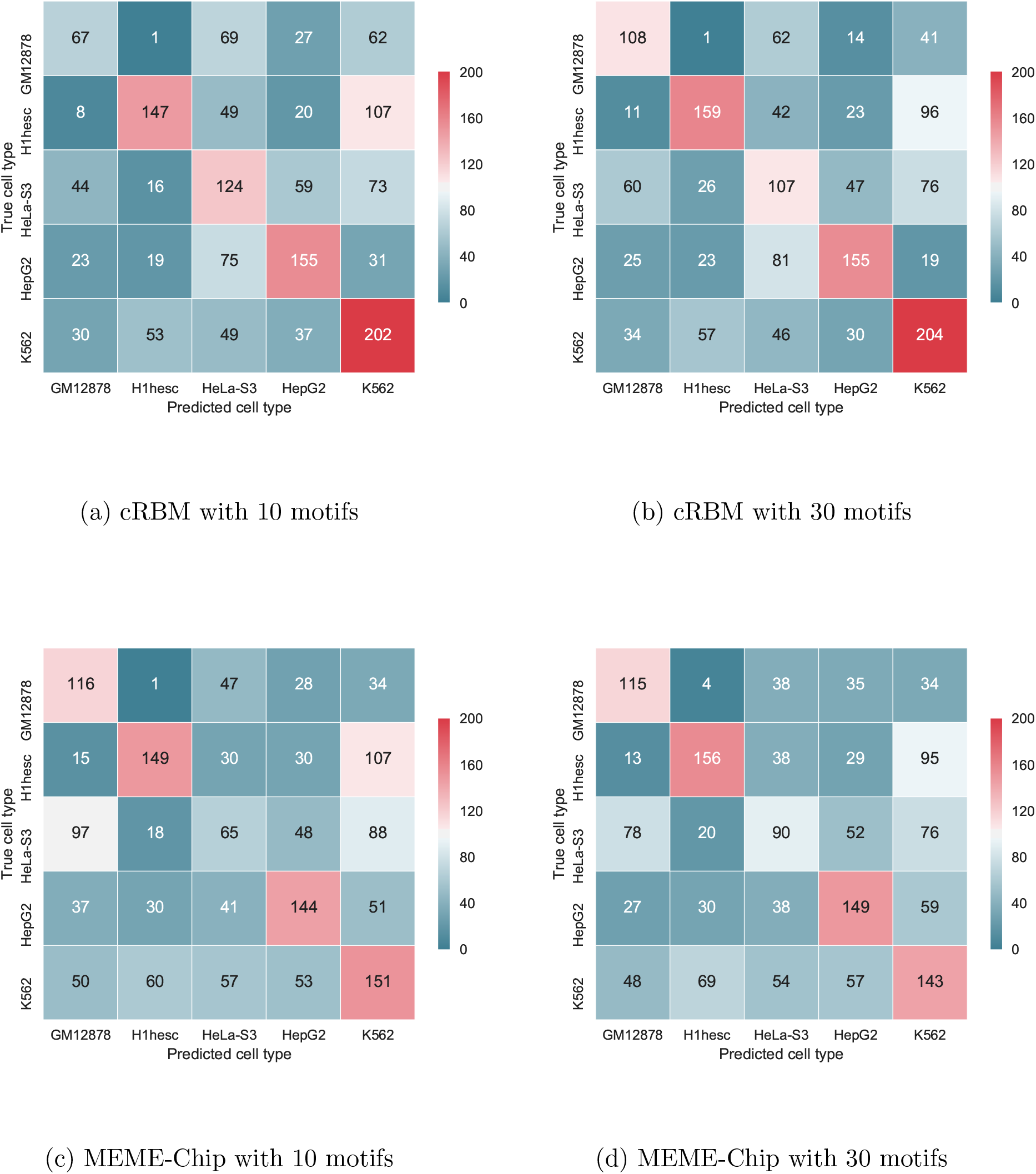
Confusion matrices showing the discriminative performance for predicting the cell-type label of *JunD* peaks. The predictions are based on cRBM (top row) and the MEME-Chip motifs, respectively (bottom row). The corresponding motifs for a) and c) are depicted in Suppl. Figure S12-S13. Motifs for b) and d) are not shown.

**Figure S15:**
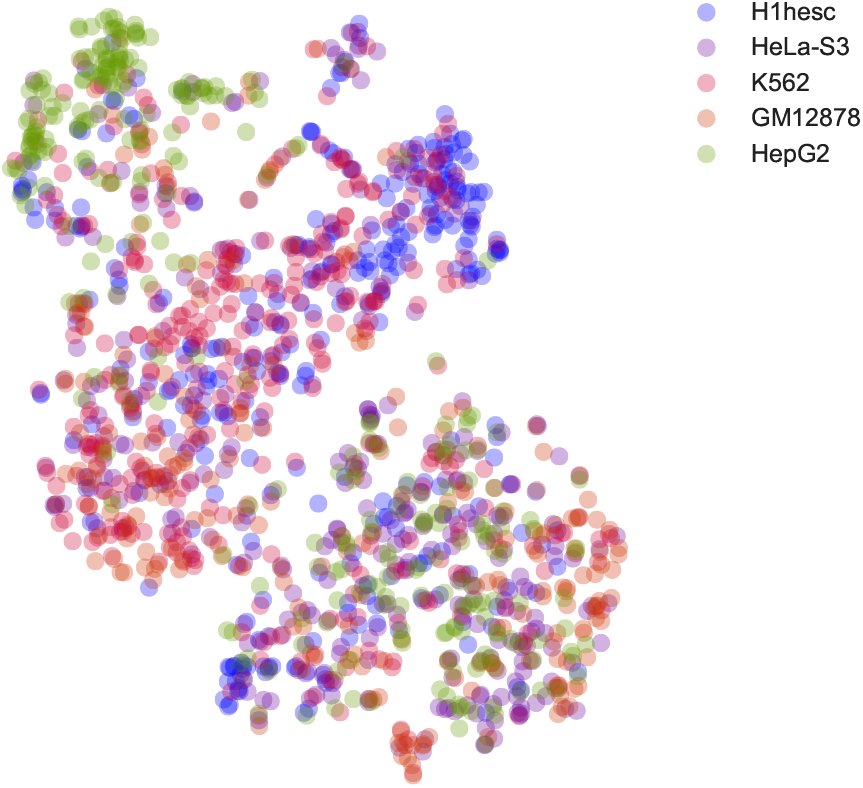
t-SNE clustering based on cRBM motif abundances of *JunD* peaks (see Suppl. Figure S12) showing all cell-types simultaneously.

## References

[1] Machanick, P. and Bailey T.L. (2011) MEME-ChIP: motif analysis of large DNA datasets. Bioinformatics, 27(12), 1696–1697.

[2] Thomas-Chollier, M., Sand, O., Turatsinze, J.V., Janky, R., Defrance, M., Vervisch, E., Brohee, S., and van Helden, J. (2008) RSAT: regulatory sequence analysis tools. Nucleic acids research, 36(suppl 2), W119–W127.

[3] Bailey, T.L., Boden, M., Buske, F.A., Frith, M., Grant, C.E., Clementi, L., Ren, J., Li, W.W., and Noble, W.S. (2009) MEME SUITE: tools for motif discovery and searching. Nucleic acids research, 37(suppl 2), W202–W208.

[4] Sandelin, A., Alkema, W., Engstro¨m, P., Wasserman, W.W., and Lenhard, B. (2004) JASPAR: an open-access database for eukaryotic transcription factor binding profiles. Nucleic acids research, 32(suppl 1), D91–D94.

[5] Wingender, E., Dietze, P., Karas, H., and Knu¨ppel, R. (1996) TRANSFAC: a database on transcription factors and their DNA binding sites. Nucleic acids research, 24(1), 238–241.

[6] Kulakovskiy, I.V., Medvedeva, Y.A., Schaefer, U., Kasianov, A.S., Vorontsov, I.E., Bajic, V.B., and Makeev, V.J. (2012) HOCOMOCO: a comprehensive collection of human transcription factor binding sites models. Nucleic acids research, 41(D1), D195–D202.

[7] Spitz, F. and Furlong, E.E. (2012) Transcription factors: from enhancer binding to developmental control. Nature reviews. Genetics, 13(9), 613.

[8] Alipanahi, B., Delong, A., Weirauch, M.T., and Frey, B.J. (2015) Predicting the sequence specificities of DNA-and RNA-binding proteins by deep learning. Nature biotechnology, 33(8), 831–838.

[9] Zhou, J. and Troyanskaya, O.G. (2015) Predicting effects of noncoding variants with deep learning–based sequence model. Nature methods, 12(10), 931.

[10] Kelley, D.R., Snoek, J., and Rinn, J.L. (2016) Basset: learning the regulatory code of the accessible genome with deep convolutional neural networks. Genome research, 26(7), 990–999.

[11] Lee, H., Grosse, R., Ranganath, R., and Ng, A.Y. (2009) Convolutional deep belief networks for scalable unsupervised learning of hierarchical representations. In Proceedings of the 26th annual international conference on machine learning, ACM pp. 609–616.

[12] Consortium, E.P. et al. (2012) An integrated encyclopedia of DNA elements in the human genome. Nature, 489(7414), 57.

[13] Hinton, G.E. A practical guide to training restricted boltzmann machines pp. 599–619 Springer Berlin Heidelberg Berlin, Heidelberg (2012).

[14] Stormo, G.D. (2000) DNA binding sites: representation and discovery. Bioinformatics, 16(1), 16–23.

[15] Maaten, L.v.d. and Hinton, G. (2008) Visualizing data using t-SNE. Journal of Machine Learning Research, 9(Nov), 2579–2605.

[16] Srivastava, P.K., Hull, R.P., Behmoaras, J., Petretto, E., and Aitman, T.J. (2013) JunD/AP1 regulatory network analysis during macrophage activation in a rat model of crescentic glomerulonephritis. BMC systems biology, 7(1), 93.

[17] Odrowaz, Z. and Sharrocks, A.D. (2012) ELK1 uses different DNA binding modes to regulate functionally distinct classes of target genes. PLoS genetics, 8(5), e1002694.

[18] Morin, J.A., Cerr´on, F., Jarillo, J., Beltran-Heredia, E., Ciesielski, G.L., Arias-Gonzalez, J.R., Kaguni, L.S., Cao, F.J., and Ibarra, B. (2017) DNA synthesis determines the binding mode of the human mitochondrial single-stranded DNA-binding protein. Nucleic Acids Research,.

[19] Tieleman, T. (2008) Training restricted Boltzmann machines using approximations to the likelihood gradient. In Proceedings of the 25th international conference on Machine learning, ACM pp. 1064–1071.

[20] Pedregosa, F., Varoquaux, G., Gramfort, A., Michel, V., Thirion, B., Grisel, O., Blondel, M., Prettenhofer, P., Weiss, R., Dubourg, V., Vanderplas, J., Passos, A., Cournapeau, D., Brucher, M., Perrot, M., and Duchesnay, E. (2011) Scikit-learn: Machine Learning in Python. Journal of Machine Learning Research, 12, 2825–2830.

[21] Theano Development Team (May, 2016) Theano: A Python framework for fast computation of mathematical expressions. arXiv e-prints, abs/1605.02688.

